# Wood capacitance is related to water content, wood density, and anatomy across 30 temperate tree species

**DOI:** 10.1101/772764

**Authors:** Kasia Ziemińska, Emily Rosa, Sean M. Gleason, N. Michele Holbrook

## Abstract

Water released from wood tissue during transpiration (capacitance) can meaningfully affect daily water use and drought response. To provide context for better understanding of capacitance mechanisms, we investigated links between capacitance and wood anatomy. On twig wood of 30 temperate angiosperm tree species, we measured capacitance, water content, wood density, and anatomical traits, i.e., vessel dimensions, tissue fractions, and vessel-tissue contact fractions (fraction of vessel circumference in contact with other tissues). Across all species, the strongest predictors of capacitance were wood density (WD) and predawn lumen volumetric water content (VWC_L-pd_, *r*^2^_adj_=0.44, *P*<0.0001). Vessel-tissue contact fractions explained an additional ∼10% of the variation in capacitance. Regression models were not improved by including predawn relative water content (RWC_pd_) or tissue lumen fractions. Among diffuse-porous species, VWC_L-pd_ and vessel-ray contact fraction were the best predictors of capacitance, whereas among ring/semi-ring-porous species, VWC_L-pd_, WD and vessel-fibre contact fraction were the best predictors. Mean RWC_pd_ was 0.65±0.13 and uncorrelated with WD. VWC_L-pd_ was weakly negatively correlated with WD. Our findings imply that capacitance depends on the amount of stored water, tissue connectivity and the bulk wood properties arising from WD (e.g., elasticity), rather than the fraction of any particular tissue.

## INTRODUCTION

Water stored in wood can buffer excessive water demand on diurnal (Goldstein *et al.* 1998; Meinzer, James & Goldstein 2004; Meinzer *et al.* 2008; Meinzer, Johnson, Lachenbruch, McCulloh & Woodruff 2009; Scholz *et al.* 2007; Köcher, Horna & Leuschner 2013; Lachenbruch & McCulloh 2014; Carrasco *et al.* 2015) and seasonal time scales (Hao, Wheeler, Holbrook & Goldstein 2013; Pratt & Jacobsen 2017; Salomón, Limousin, Ourcival, Rodríguez-Calcerrada & Steppe 2017). Estimates of the contribution of stored water to a tree’s daily water budget range from 5 to 50% (Goldstein *et al.* 1998; Kobayashi & Tanaka 2001; Phillips *et al.* 2003; Meinzer *et al.* 2004; Scholz *et al.* 2007; Köcher *et al.* 2013; Carrasco *et al.* 2015). Thus, water storage can be an important component of whole-plant water balance (Gleason, Blackman, Cook, Laws & Westoby 2014; Blackman *et al.* 2016; Christoffersen *et al.* 2016). However, the mechanisms of water storage and release, and their structural underpinnings remain unclear. Consequently, our understanding of cost vs. benefits of water storage and how storage is coordinated with other tree functions is limited. Here, our objective was to quantify water storage (amount of stored water) and diurnal capacitance (water released per wood volume per change in stem water potential, kg m^−3^ MPa^−1^) across a diverse suite of 30 temperate angiosperm trees.

Capacitance is most often measured using psychrometers and estimated as the initial slope extracted from a water release-potential curve (Meinzer, James, Goldstein & Woodruff 2003; Meinzer *et al.* 2008; Scholz *et al.* 2007; McCulloh *et al.* 2012; Trifilò *et al.* 2015; Jupa, Plavcová, Gloser & Jansen 2016; Santiago *et al.* 2018; Siddiq, Zhang, Zhu & Cao 2019). However, if tissues *in natura* never become fully saturated, or if the operating water potential of xylem falls outside the initial water release curve, then this way of estimating capacitance may be functionally irrelevant. Oversaturating wood prior to measuring capacitance may also skew results. Moreover, using psychrometers on excised material is likely burdened with an artefact due to water being released from open vessel ends (Tyree & Yang 1990; Jupa *et al.* 2016). Few studies have estimated diurnal capacitance across the range of water potentials experienced by field plants, with only two studies using pressure chamber measurements of non-transpiring leaves (Zhang, Meinzer, Qi, Goldstein & Cao 2013; Wolfe & Kursar 2015) and one study using psychrometers on excised tissues (Richards, Wright, Lenz & Zanne 2014). We addressed this shortcoming by measuring capacitance between predawn and midday during peak summer conditions. To avoid open vessels and oversaturation artefacts, we estimated capacitance from the difference in wood water content between predawn and midday with the corresponding change in stem water potential measured using bagged, non-transpiring (equilibrated) leaves (Begg & Turner 1970; Clearwater & Meinzer 2001). We refer to this measure as ‘day capacitance’, following Zhang *et al.* (2013).

Anatomical structure should determine wood capacitance (Zimmermann 1983; Tyree & Yang 1990; Holbrook 1995). Tyree & Yang (1990) proposed that water release consists of three phases linked to anatomy: 1) an initial phase (0 to *ca*. −0.6 MPa), when water is released from capillary storage (already embolized fibres, vessels and tracheids, and intercellular spaces), 2) a second phase (< −0.6 MPa but prior to vessel embolization), when water is released from elastic storage (cells with elastic cell walls), and 3) a final phase (below the embolization threshold), when water is released from embolizing vessels and fibres, likely resulting in permeant damage to the xylem tissue, at least in large trees. Despite this well-developed and popular theory, few studies have quantified wood anatomy in relation to capacitance, and none of them used methods, which would avoid open vessels and oversaturation artefacts. As a result, we lack quantitative understanding of the anatomical drivers of capacitance, especially for angiosperm trees, which exhibit substantial variation in stem cell sizes, types, their proportions and geometry.

Given these knowledge gaps, it is surprising that parenchyma, the main living tissue in wood commonly believed to have elastic cell walls, is often assumed the primary source of capacitance in stems (e.g, Meinzer *et al.* 2003; Steppe & Lemeur 2007; Plavcová & Jansen 2015; Vergeynst, Dierick, Bogaerts, Cnudde & Steppe 2015; Morris *et al.* 2016; Li *et al.* 2018; Rungwattana & Hietz 2018; Santiago *et al.* 2018). Several studies have extended this idea to suggest that a higher parenchyma fraction may therefor confer higher capacitance (Borchert & Pockman 2005; Scholz, Phillips, Bucci, Meinzer & Goldstein 2011; Pratt & Jacobsen 2017; Secchi, Pagliarani & Zwieniecki 2017; Nardini, Savi, Trifilò & Lo Gullo 2018). While this may indeed be the case in highly parenchymatous stems like palms or baobabs (Holbrook & Sinclair 1992; Chapotin, Razanameharizaka & Holbrook 2006a b), it may be less likely for the majority of tree species, which tend to have lower wood parenchyma fractions (Morris *et al.* 2016). In these species, parenchyma are connected with other cells (fibres, vessels) via highly lignified middle lamella impeding parenchyma independent volumetric changes. Moreover, recent microCT studies, suggest that water released from cavitating fibres and vessels may contribute meaningfully to capacitance at moderate water potentials (Knipfer *et al.* 2017, 2019).

Here, we measured detailed anatomical traits, water storage and capacitance to address the following questions: 1) how large is day capacitance and how does it vary among diverse temperate tree species?, 2) how much water is stored in wood and does this water volume limit day capacitance during peak summer?, 3) which anatomical traits are related to capacitance?

## MATERIALS AND METHODS

### Site, species and individuals

All measurements were taken from trees growing at the Arnold Arboretum of Harvard University in Boston (Massachusetts, USA). Mean annual temperature in 2017, the year when capacitance measurements were taken, was 10.9°C and annual precipitation was 787 mm. Growing season (March to October) mean temperature was 15.3°C and precipitation was 562 mm. Mean temperature during sampling (August) was 21.2°C and precipitation was 22 mm. The soil texture is variable across the Arboretum and sampled trees grew on: sandy loam, slit loam, loamy sand, and an outcrop complex. The studied trees were excluded from the Arboretum’s supplemental watering regime during the summer 2017.

Thirty species of deciduous, angiosperm trees were selected, spanning 30 genera and 26 families (Table 1; tree accession numbers are listed in Table S1). InsideWood database (InsideWood 2004; Wheeler 2011) was used as a guide to select species with the most diverse anatomies, e.g., parenchyma abundance, porosity (diffuse-, ring- or semi-ring-porous) and vessel size.

**Table 1.**
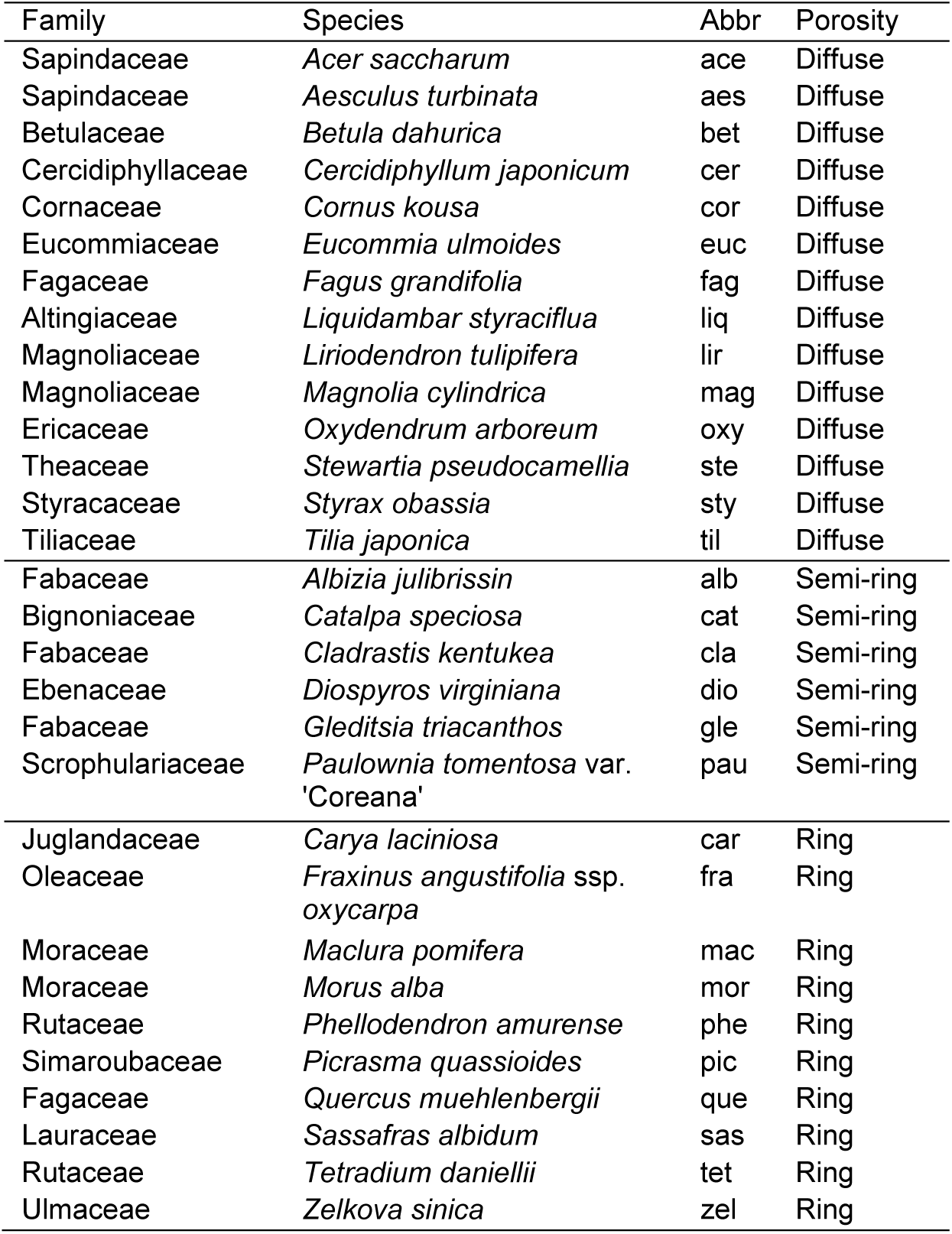
Species list, their abbreviations and porosity type.

Due to the limited number of individuals per species, trees were sampled across the existing variation in topography and soil texture. Trees growing near streams and ponds were avoided and only healthy trees were sampled. When possible, similar height trees per given species were studied. We measured branches exposed to the midday sun. However, variation in forest density near the measured trees contributed to variation in the amount of sun branches received throughout the course of a day and across measurements and species.

### Capacitance and water storage: sampling and measurements

Water storage and capacitance measurements were performed during the second half of August 2017. Ten individuals per day were sampled, resulting in a total of nine sampling days. In the case of rain, sampling was delayed for one or two days. For stem water potential, four leaves per tree were enclosed in opaque, silver, 4 mil (0.1 mm) thick Mylar zip-lock bags the evening before sampling. In principle, the water potential of a bagged, non-transpiring leaf will come into equilibrium with the stem water potential. As such, bagged leaf water potential (i.e., after equilibration) provides an estimate of stem water potential (Begg & Turner 1970; Clearwater & Meinzer 2001). Two leaves at predawn and two leaves at midday were collected. The bagged leaf was cut with a razor blade and the bag was immediately closed and placed in an insulated box at ambient temperature. Six terminal, 0.5-0.7 m long twigs per tree, four of which also included the leaves bagged the previous evening, were sampled. Three of these twigs (per tree) were collected at predawn and three twigs at midday. Twigs for each predawn-midday pair were located near each other, as close as tree architecture allowed (usually within 0.5 m) to limit the variation in water content that might be expected along the length of a branch. After collecting the predawn twig, the cut surface on the branch still attached to a tree was covered with Vaseline to reduce drying from the exposed surface. Twigs were quickly de-leafed (apical meristem was removed together with most distal leaves), double-bagged in zip-lock bags, and placed in an additional opaque bag at ambient temperature. Each sampling event took approximately two hours. After collecting all leaves and twigs, material was transported to the onsite laboratory within 5 min.

Immediately after arrival to the laboratory, leaf water potential was measured on one leaf per tree using a pressure chamber (Model 1000, PMS Instrument Company, USA). Next, twig segments *ca*. 60 mm long and 5 mm diameter (excluding bark and pith) were cut at a minimum 50 mm distance from the initial, field cut. The segments were quickly wrapped in Parafilm, placed in a 4 mil zip-lock bag, and then into a cooled box with ice. After preparing all samples, the following steps were carried out in a humidity-controlled room (at relative humidity of 75-80%) to minimize water loss. Bark and pith were removed, the ends were trimmed by a few mm, and fresh mass was measured on an analytical balance (Sartorius, 0.00001g), after which the samples were placed in distilled water.

These wood samples were then stored at 4°C for two weeks and, afterwards, saturated mass and volume was measured using Archimedes principle as described in (Zieminska, Westoby & Wright 2015). In several cases, twigs did not sink even after two weeks of soaking, thus indicating that gas was still present in those samples. These samples were held under water at *ca*. 60°C for several hours. This treatment resulted, in all cases, in samples that sank, and, next, volume and saturated mass was recorded. For logistic reasons, we were unable to measure volume within 48 hours after sampling, as is commonly done in other studies. However, our preliminary tests confirmed that across all species studied, volume measured on fresh vs. saturated samples (after two weeks of soaking at 4°C) differed on average by 2% ± 1.1% SD, and as such, should not meaningfully influence our volume estimates. As comparison, soaking for 48 hours resulted in 1.6% ± 0.9% SD volume change – similar to the change after two weeks. After saturated mass and volume measurements, the samples were dried at 102°C for three days and dry mass was recorded.

Broadly, water storage is defined here as the amount of water contained in a wood sample. It can be expressed as: relative water content or volumetric water content. Relative water content (RWC) is the proportion of water in a sample relative to the maximum amount of water that could be stored in that sample and was calculated as follows:

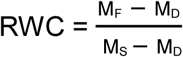

where M_F_ is sample fresh mass, M_D_ is sample dry mass, and M_S_ is sample saturated mass. Volumetric water content (VWC) indicates total water volume per sample volume (Gartner, Moore & Gardiner 2004) and was calculated as follows:

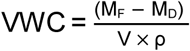

where V is sample volume and ρ is water density, assumed to equal 1 g cm^−3^. We also partitioned VWC to lumen (VWC_L_) and wall (VWC_W_) water content. For that, we assumed that the fibre saturated point (FSP), defined as the point where only water bound in cell wall is present, is at 30% moisture content (MC) (Ross 2010; Dlouhá, Alméras, Beauchêne, Clair & Fournier 2018). The standard equation for moisture content is:

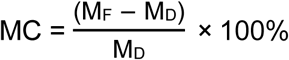

(Ross 2010). From this equation, we calculated sample mass at FSP (M_FSP_) and estimated VWC_L_ as:

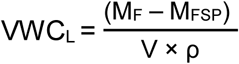

and VWC_W_ as:

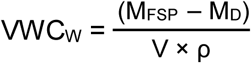

Next, we estimated lumen relative water content (RWC_L_, Longuetaud *et al.* 2016):

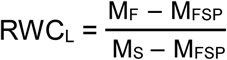

All water content indices were estimated at predawn and midday, indicated by subscripts (_pd_ and _md_).

Following a simplified version of the equation from Meinzer *et al.* (2003) and Richards *et al.* (2014), cumulative water released (CWR, kg m^−3^) was calculated, separately for predawn and midday, as follows:

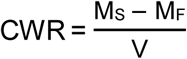

and then multiplied by 1000 to convert units from g cm^−3^ to kg m^−3^. Wood day capacitance (kg m^−3^ MPa^−1^) was estimated as:

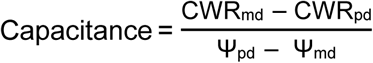

where Ψ is stem water potential. This calculation of capacitance takes saturation as a reference point, to minimize the effect of intrinsic differences between predawn and midday samples (e.g., in their wood density). However, capacitance could also be estimated more directly from fresh and dry samples only:

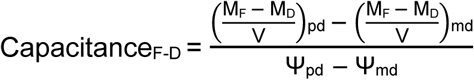

In principle, if predawn and midday water content was measured on exactly the same samples, the two measures of capacitance would lead to exactly same results. However, this was not the case in our study, because predawn and midday water content was assessed on separate samples. Nevertheless, the two measures were well correlated with each other (*r*^2^=0.76, *P*<0.001) and the intercept and slope were not significantly different from 0 and 1, respectively (P=0.80 and P=0.59; Fig. S1 in supporting information; *Paulownia tomentosa* was excluded because, as an extreme outlier (see Results), it would have a disproportionately strong effect on this analysis). This strong correlation suggests that findings based on ‘capacitance’ as presented in the current study would be very similar to findings based on ‘capacitance_F-D_’.

All measured traits, their abbreviations used in the text and units are listed in Table 2.

**Table 2.** Traits, abbreviations and units. Fractions of wall (w), lumen (l) and wall+lumen (w+l) of a given tissue were measured. Subscripts denote: lumen (L), predawn (pd), midday (md) and difference between predawn and midday (pd-md).

| Trait | Abbr | Units |
| --- | --- | --- |
| Day capacitance | capacitance | $\text{kg m}^{-3} \text{MPa}^{-1}$ |
| Cumulative water released | $\text{CWR}_{\text{pd}}, \text{CWR}_{\text{md}}, \Delta\text{CWR}_{\text{md-pd}}$ | $\text{kg m}^{-3}$ |
| Stem water potential | $\Psi_{\text{pd}}, \Psi_{\text{md}}, \Delta\Psi_{\text{pd-md}}$ | MPa |
| Volumetric water content | $\text{VWC}_{\text{pd}}, \text{VWC}_{\text{md}}, \Delta\text{VWC}_{\text{pd-md}}$ | Unitless |
| Volumetric water content: saturated | $\text{VWC}_{\text{sat}}$ | " |
| Lumen volumetric water content | $\text{VWC}_{\text{L-pd}}, \text{VWC}_{\text{L-md}}, \Delta\text{VWC}_{\text{L-pd-md}}$ | " |
| Lumen volumetric water content: saturated | $\text{VWC}_{\text{L-sat}}$ | " |
| Wall volumetric water content | $\text{VWC}_{\text{w}}$ | " |
| Relative water content | $\text{RWC}_{\text{pd}}, \text{RWC}_{\text{md}}, \Delta\text{RWC}_{\text{pd-md}}$ | " |
| Lumen relative water content | $\text{RWC}_{\text{L-pd}}, \text{RWC}_{\text{L-md}}, \Delta\text{RWC}_{\text{L-pd-md}}$ | " |
| Wood density | WD | $\text{g cm}^{-3}$ |
| Fibre fraction: w, l, w+l |  | Unitless |
| Axial+ray parenchyma fraction: w, l, w+l |  | " |
| Axial parenchyma fraction: w, l, w+l |  | " |
| Ray fraction: w, l, w+l |  | " |
| Vessel fraction: w, l, w+l |  | " |
| Living fibre fraction: w, l, w+l |  | " |
| Conduits<sub>15</sub> fraction: w+l |  | " |
| Vessel-fibre contact fraction |  | " |
| Vessel-axial+ray parenchyma contact fraction |  | " |
| Vessel-axial parenchyma contact fraction |  | " |
| Vessel-ray contact fraction |  | " |
| Vessel-vessel contact fraction |  | " |
| Vessel-conduits<sub>15</sub> contact fraction |  | " |
| Vessel mean area | | $\mu\text{m}^2$ |
| Vessel mean diameter | | $\mu\text{m}$ |
| Hydraulically weighted vessel diameter | $D_{\text{H}}$ | " |
| Vessel number per area | | $\text{mm}^{-2}$ |
| Vessel size to number ratio | | $\text{mm}^2 \text{mm}^{-2}$ |
| Lumen fraction per total fibre area | $\text{lumen fraction}_{\text{fibre}}$ | Unitless |
| Lumen fraction per total axial+ray parenchyma area | $\text{lumen fraction}_{\text{axial+ray}}$ | " |
| Lumen fraction per total axial parenchyma area | $\text{lumen fraction}_{\text{axial}}$ | " |
| Lumen fraction per total rays area | $\text{lumen fraction}_{\text{ray}}$ | " |
| Lumen fraction per total vessels area | $\text{lumen fraction}_{\text{vessel}}$ | " |
| Lumen fraction per total living fibres area | $\text{lumen fraction}_{\text{living fibre}}$ | " |

### Anatomy: sampling and measurements

For anatomical measurements, one, midday sun-exposed twig per tree was collected in August 2016. Twigs were transported to the lab in semi-opaque, plastic bags. Next, several pieces with wood diameter of 4-6mm (excluding bark and pith) were cut and stored in 70% ethanol for about three months until further processing. In about a third of cases, samples collected in 2016 and 2017 came from different individuals (Table S1) due to either deteriorated foliage health state between the two years or considerable storm damage. Twig size (diameter and length) and location on the tree (e.g., aspect) were consistent between 2016 and 2017 sampling, although in some cases slightly lower branches were measured in 2017 to allow for leaf bagging.

Cross-sections ∼10-20 µm thick were made using Reichert sledge microtome, blade holder (Accu-Edge M7321-43) and low-profile blades (Accu-Edge, 4689). To facilitate flattening, sections were stored between two glass slides held together by paper clips in 50% ethanol for several days. Next, sections were stained in a mixture of safranin O and Alcian Blue (0.35g safranin O in 35ml 50% ethanol + 0.65g of Alcian Blue in 65ml of distilled water) for three minutes, and then rinsed and mounted on a glass slide in glycerol. Longitudinal radial and tangential sections of one sample per species were also taken to assist in anatomical interpretations. A pie-shaped region of one cross-section per tree (three trees per species) was photographed using a Zeiss Axiophot microscope, AxioCam 512 camera and Zen Blue 2.3 software (ZEISS, Germany). Magnification was provided by a Plan-Neofluar 20x objective. Several photos were taken to cover the entire pie region, stretching from pith to cambium, which then were stitched together in freeware Image Composite Editor (Microsoft, 2015).

Anatomical measurements were done on three individuals per species (Table S1), across all growth rings (4.4 ± 1.5 SD) with the exception of vessel-tissue contact fractions (see below), which were measured on all except the innermost ring due to the substantial time/labour cost of this measurement. The potential error resulting from this approach is likely minimal because tissue fractions measured on all growth rings exhibited a strong correlation with tissue fractions measured on all except innermost ring (*r*^2^>0.90 for most tissues, *r*^2^= 0.79 for fibre fraction and *r*^2^= 0.76 for living fibre fraction), and the innermost ring usually contributes little to the whole-twig cross-section.

Lumen and wall were measured separately for all tissue fractions: fibre, living fibre, axial parenchyma, ray parenchyma, vessels, and conduits with maximum diameter <15 µm (for these, lumen and wall were counted together). The latter category (denoted hereafter conduits_15_) likely encompassed small vessels, the tapered ends of vessels (tails), and tracheids. Living fibres were identified by the presence of starch, nuclei or septa, and their wall thickness was similar to fibres and/or thicker than parenchyma walls. From these measurements, we calculated the fraction of lumen per given tissue. The vessel characteristics measured included: vessel mean area and diameter (mean of minimum and maximum diameters of a given vessel), hydraulically weighed diameter (D_H_ = Σdiameter^5^ / Σdiameter^4^; Sperry, Nichols, Sullivan & Eastlack 1994), vessel number per cross-sectional area, and the vessel size-to-number ratio (denoted S in Zanne *et al.* 2010), calculated as mean vessel area divided by vessel number per cross-sectional area). We also measured the proportion of vessel circumference in contact with other tissues, referred to collectively as ‘vessel-tissue contact fraction’. For example, ‘vessel-axial parenchyma contact fraction’ is the proportion of vessel circumference in contact with axial parenchyma.

For vessel traits, we first colored vessels in black using the “magic wand” tool in freeware GIMP (GNU Image Manipulation Program, www.gimp.org) excluding protoxylem vessels. Next, vessels and the total pie-shaped sample area were automatically measured in freeware ImageJ (ImageJ; Schneider, Rasband & Eliceiri 2012). Image manipulation and analysis were processed using Wacom Cintiq 22HD pen display (Wacom Technology Corporation, Portland, USA).

For tissue fractions, a grid method was applied (Zieminska *et al.* 2015). Briefly, each grid point overlaid over the pie region in ImageJ was classified (‘Cell counter’ plugin, https://imagej.net/Cell_Counter) depending on which tissue it fell into (Fig. 1 in Zieminska *et al.* 2015). The distance between grid points was 55 µm. On average, we analysed 484 (±128) points per pie region and the total number of points depended on the wood sample and the pie region sizes. For ray wall fraction we counted only walls perpendicular to the cross-section, and as such our measurements are likely underestimates. For vessel-tissue contact fractions, all vessels in a pie region were analysed except for the innermost growth ring (see above). For this measurement, we modified the grid method. In Photoshop CS4 (Adobe Systems Incorporated, USA), dots were placed at equal distances to each other at 70 pixels (22.5 µm, Fig. 1) along the vessel lumen circumference, with the exception of the first and last dot, whose distance to each other was determined by vessel circumference. We counted the last dot only if its distance from the first dot was larger than 35 pixels (half-distance between all other dots). Next, using the ‘Cell counter’ plugin as for tissue fractions, we classified each dot depending on the neighbouring tissue. For instance, if the dot fell on the border with ray, it was classified as vessel-ray contact dot, and so forth. We then took the ratio of all vessel-ray contact dots to total analysed dots in a pie region, and did the same for the other tissues. This ‘dot-ratio’ is vessel-tissue contact fraction for a given, abutting tissue. On average, we analysed 943 dots (±605) per each pie region. The number of dots depended on the total vessel circumference (e.g., it was the highest for species with large vessel fraction composed of many small vessels).

**Figure 1.**
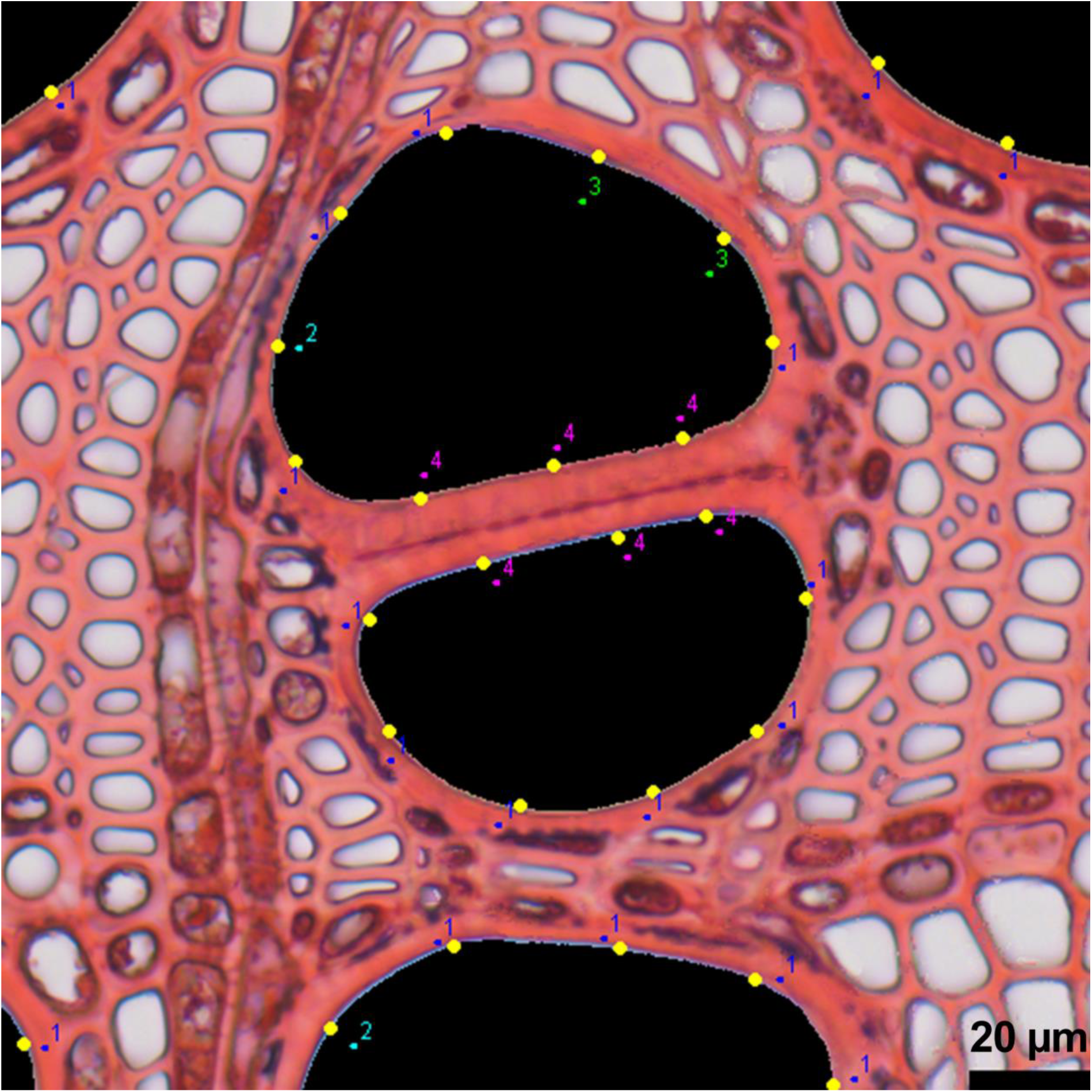
Illustration of the method used to estimate vessel-tissuecontact fractions between vessels and other tissues (fibres, axial parenchyma, rays, vessels, conduits_15_). The image shows fragment of a cross-section of *Fraxinus angustifolia* ssp. *oxycarpa*. Yellow dots distributed on vessel (black) circumference are classified based on the tissue in contact with a vessel (1 – axial parenchyma, 2 – rays, 3 – fibres, 4 – vessels; conduits_15_ – not shown).

### Wood density

Wood density (WD) was measured on the same samples as water storage and capacitance measurements, as sample dry mass divided by saturated volume (g cm^−3^).

### Data analysis

The studied species encompassed 14 diffuse-porous species, 10 ring-porous species, and 6 semi-ring-porous. Anatomically, diffuse-porous species were very different from the two other groups, which in turn were very similar to each other (see Results). Consequently, we ran analyses across all species, as well as within the two porosity groups: diffuse-porous and ring/semi-ring-porous.

All analysis were performed in R (R Core Team 2018). We used linear multiple regression models to assess predictors of capacitance using the ‘lm’ function. The distribution of residuals was evaluated using the ‘residualPlot’ and ‘qqPlot’ functions in the “car” package (Fox & Weisberg 2011). Colinearity of predictors was assessed using the variance inflation factor (VIF, ‘vif’ function). When VIF>4, we removed the variables from the model. *Paulownia tomentosa* was an outlier in many relationships, so we excluded it from several models and indicate that accordingly in the Results. Correlation matrices were obtained using the ‘corrplot’ function in “corrplot” package (Wei & Simko 2017), and scatterplot matrices were obtained using the base function ‘pairs’. Selected figures (e.g., correlation matrices) were additionally edited in Illustrator (Adobe Systems Incorporated, USA).

We note that tissue fractions and vessel-tissue contact fractions are not independent, i.e., two fractions taken from the same whole should be expected to vary inversely with one another. However, this expectation is weakened when more than two tissues create the whole, as is the case for our study. Nevertheless, *r*^2^ and *P* values calculated from plotting one fraction against another should be interpreted with caution and in the context of all other tissue fractions.

To assess the relationship between total lumen fraction and saturated VWC_L_ (VWC_L-sat_), we fitted major axis models and tested for differences in slope and elevation using the ‘ma’ function in the “smatr” package (Warton, Duursma, Falster & Taskinen 2011).

## RESULTS

### Anatomical variation

Anatomical variation in tissue fractions, vessel properties and vessel-tissue contact fractions was considerable (Table S2, Fig.2a). Fibres represented the highest fraction (0.45 ± 0.08, here and thereafter we report mean ± one standard deviation), followed by axial+ray parenchyma (0.28 ± 0.08), and then vessels (0.24 ± 0.11). Average axial+ray parenchyma fraction was higher than that found by global analysis of ∼400 temperate species (0.21 ± 7.9) and ranged from 0.16 to 0.43, spanning more than half of the reported in that global analysis (Morris *et al.* 2016). Vessel fraction encompassed almost the entire spectrum of global variation (Zanne *et al.* 2010). Conduits_15_ were the least abundant (0.016 ± 0.023, absent in *P. tomentosa*). Living fibres were observed in eight species, but comprised only a small fraction of total wood area (0.026 ± 0.037) with *Acer saccharum* as an outlier (0.11). Axial and ray parenchyma had similar fractions (axial: 0.13 ± 0.09, ray: 0.15 ± 0.04). The fraction of tissue lumen per total area of a given tissue, called here ‘lumen fraction_tissue_’ (Table 2) varied considerably across tissue types, with vessels exhibiting the largest lumen fraction_vessel_ (0.82 ± 0.04), followed by rays (0.64 ± 0.05), axial parenchyma (0.52 ± 0.10), and then fibres (0.30 ± 0.15). Although fibres had the lowest lumen fraction_fibre_, it also varied the most (8-fold) across species. As mentioned in Materials and Methods, ray lumen fractions (per entire wood cross-section as well as per ray tissue) were likely overestimated (wall was likely underestimated).

**Figure 2.**
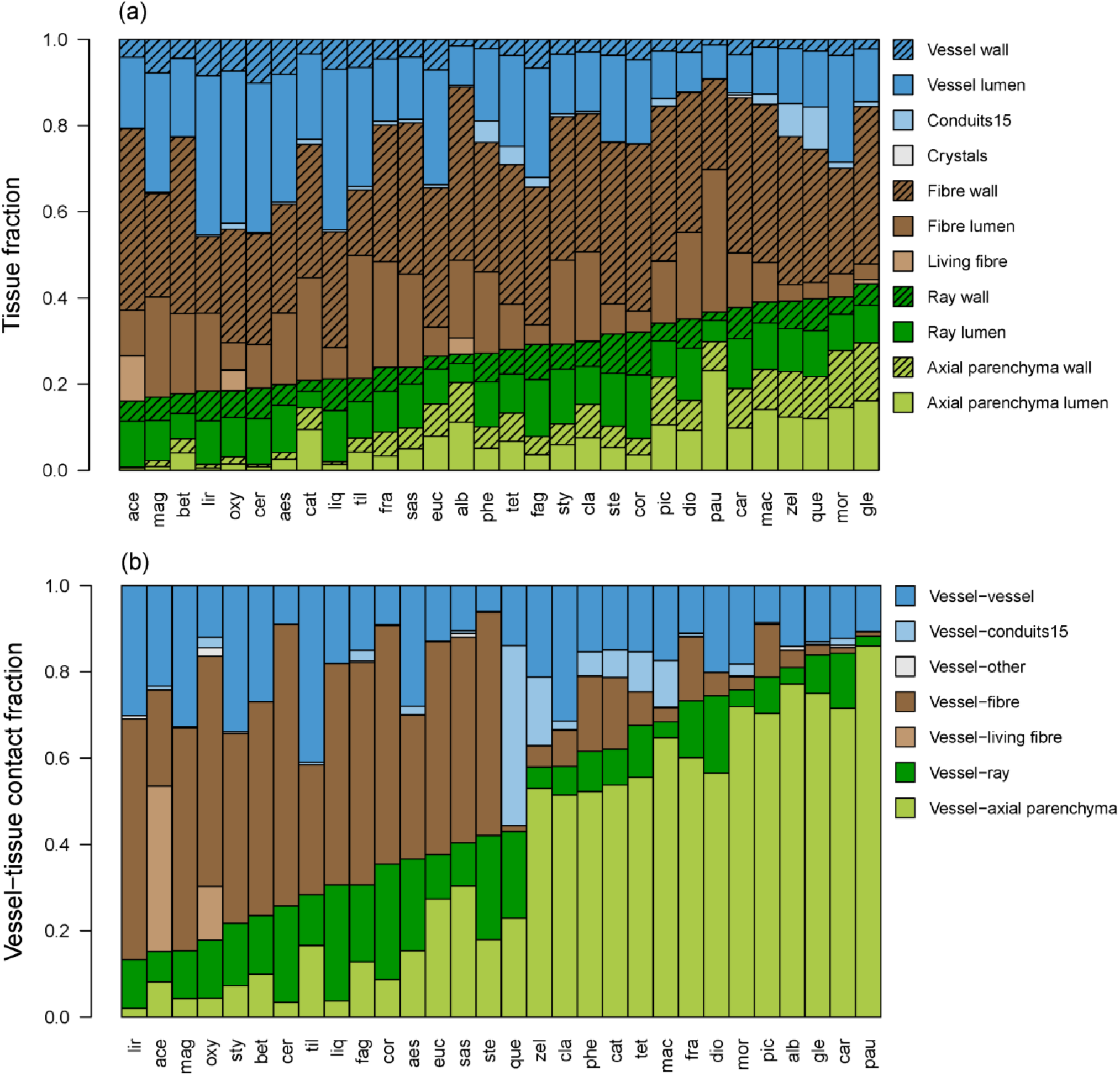
Tissue fractions sorted along axial+ray parenchyma fraction (a) and vessel-tissue contact fractions sorted along vessel-axial+ray parenchyma contact fraction (b). Three-letter codes denote species (Table 1).

Vessel-tissue contact fractions differed significantly between tissues and across species, more so than tissue fractions (Table S2, Fig.2b). The largest contact fraction was between vessels and axial parenchyma (0.36 ± 0.28) and between vessels and fibres (0.27 ± 0.22 SD). Vessel-vessel contact fraction was on average 0.18 ± 0.09, followed by vessel-ray contact (0.13 ± 0.07).

Anatomical characteristics differed considerably between diffuse-porous, semi-ring-porous and ring-porous species (Fig. 3), and these differences strongly influenced the investigated trait-trait relationships. Semi-ring-porous and ring-porous species tended to group with each other and many of these species occur in both forms (InsideWood 2004), hence we grouped them into one category denoted ‘ring/semi-ring-porous’ (Fig. 3).

**Figure 3.**
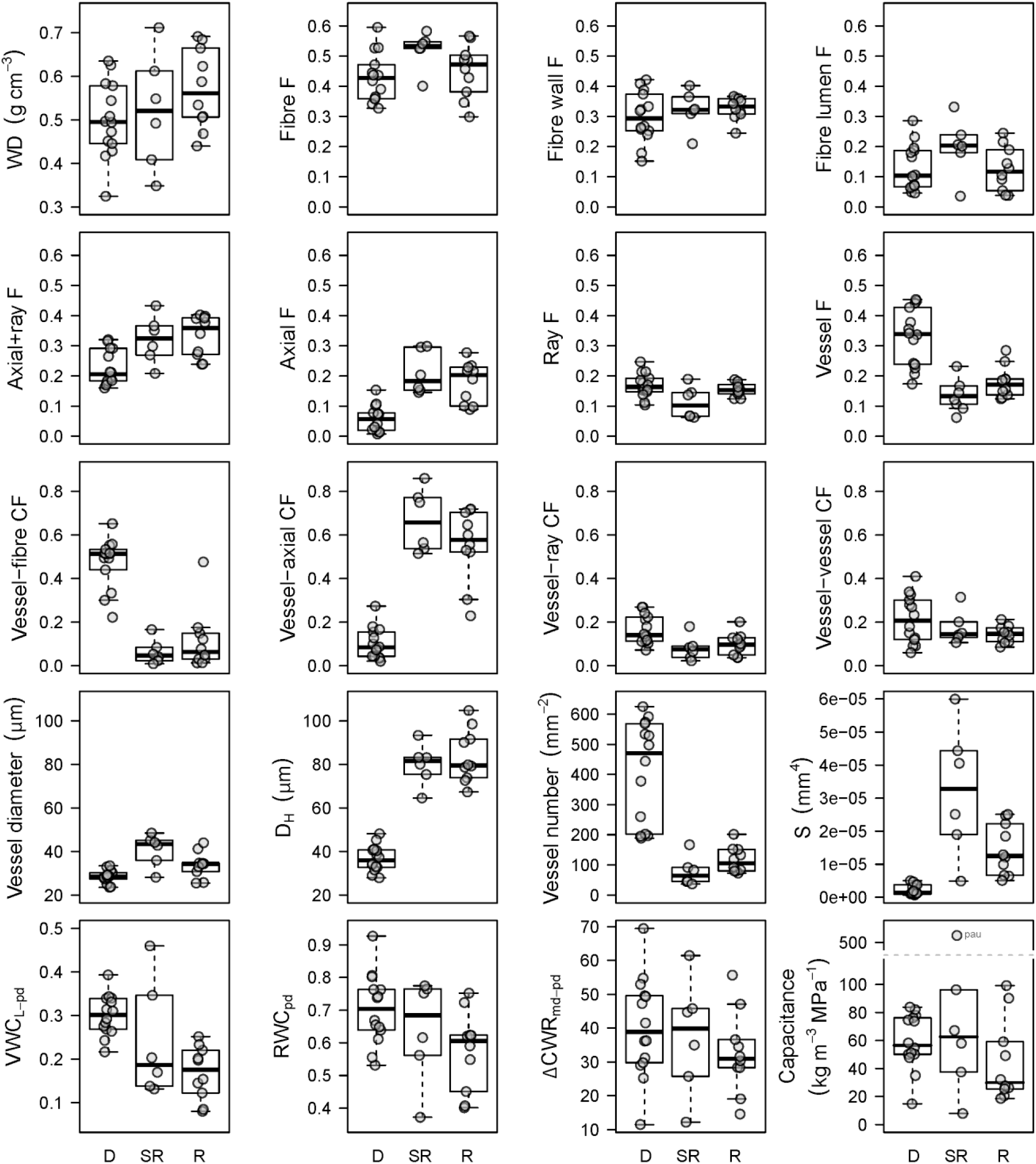
Boxplots illustrating differences between diffuse-porous (D, n=14), semi-ring-porous (SR, n=6) and ring-porous species (R, n=10). Circles denote species averages. Traits shown: wood density (WD), tissue fractions (F), vessel-tissue contact fractions (CF), hydraulically weighted diameter (D_H_), vessel size to number ratio (S), predawn lumen volumetric water content (VWC_L-pd_), predawn relative water content (RWC_pd_), midday to predawn difference in cumulative water released (ΔCWR_md-pd_) and capacitance. An outlier is *Paulownia tomentosa* (pau).

Within diffuse-porous species, fibre fraction was inversely related to vessel fraction (*r*^2^=0.68, *P*<0.001 *P*<0.001, Fig.4a), but not to axial+ray parenchyma fraction (*r*^2^=0.00, *P*=0.87, Fig. 4b). In contrast, within ring/semi-ring-porous species, fibre fraction was negatively correlated with axial+ray parenchyma fraction (*r*^2^=0.50, *P*<0.01, Fig. 4b) but not with vessel fraction (*r*^2^=0.14, *P*=0.15, Fig. 4a). Across all species, vessel fraction was negatively related to axial+ray parenchyma fraction (*r*^2^=0.43, *P*<0.001, Fig. 4c) and this was driven by porosity type, i.e., diffuse-porous species tended to have higher vessel fraction and lower axial+ray parenchyma as oppose to ring/semi-ring-porous species. Axial parenchyma fraction was negatively correlated (weakly) with ray fraction, with diffuse-porous species tending towards higher ray fraction and ring/semi-ring-porous toward higher axial parenchyma fraction (*r*^2^=0.15, *P*<0.05, Fig. S2).

**Figure 4.**
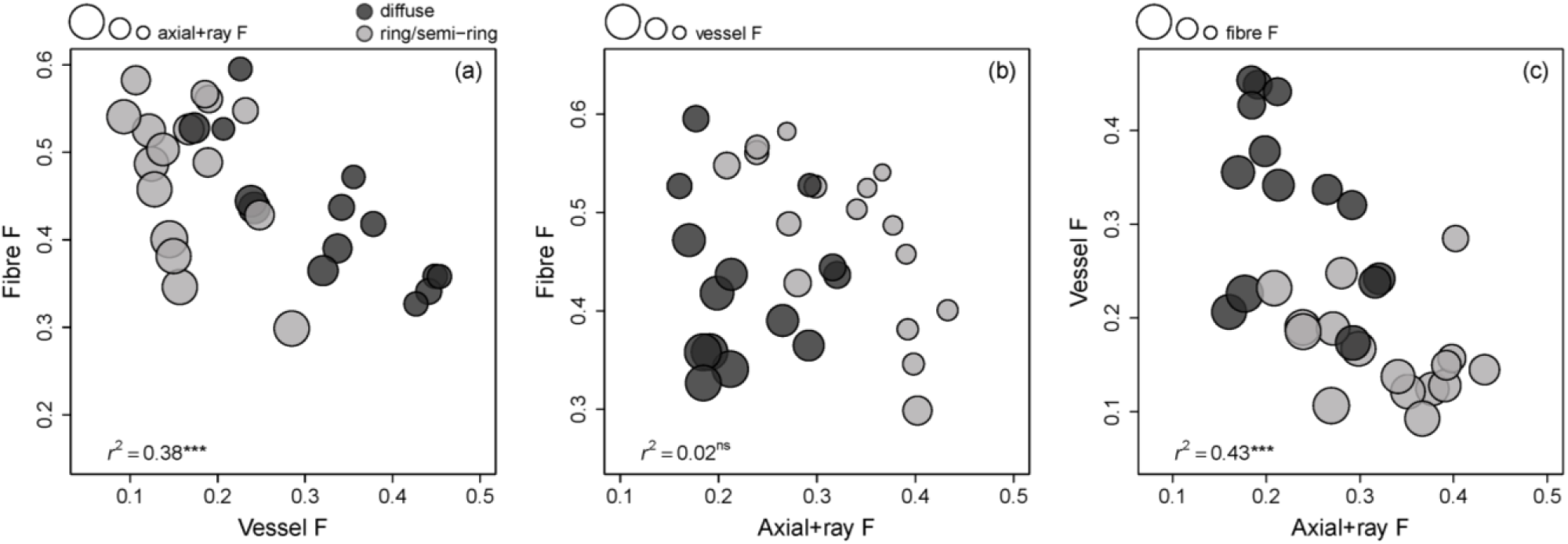
Relationships between tissue fractions (F): fibre fraction vs. vessel fraction (a), fibre fraction vs. axial+ray parenchyma fraction (b), and vessel fraction vs. axial+ray parenchyma fraction (c). Legends illustrate trait used to scale bubble size and porosity type.

Species with more abundant axial and ray parenchyma tended to have more contact between these tissues and vessels. However, having abundant fibre or vessel fractions did not translate to higher vessel-fibre or vessel-vessel contact (Fig. S2). Vessel-fibre contact fraction was inversely correlated with vessel-axial parenchyma contact fraction (*r*^2^=0.75, *P*<0.001, slope: −1.1, Fig. 5), Vessel-ray contact fraction was positively related to vessel-fibre contact fraction (*r*^2^=0.34, *P*<0.001, slope: 1.9) and negatively to vessel-axial parenchyma contact fraction (*r*^2^=0.40, *P*<0.001, slope: −2.6). The relatively steep slopes of these relationships indicated that per one unit change in vessel-ray contact fraction, the two other tissues changed by approximately two units. All these trade-offs were driven by strong differences between porosity groups (Fig. 3).

**Figure 5.**
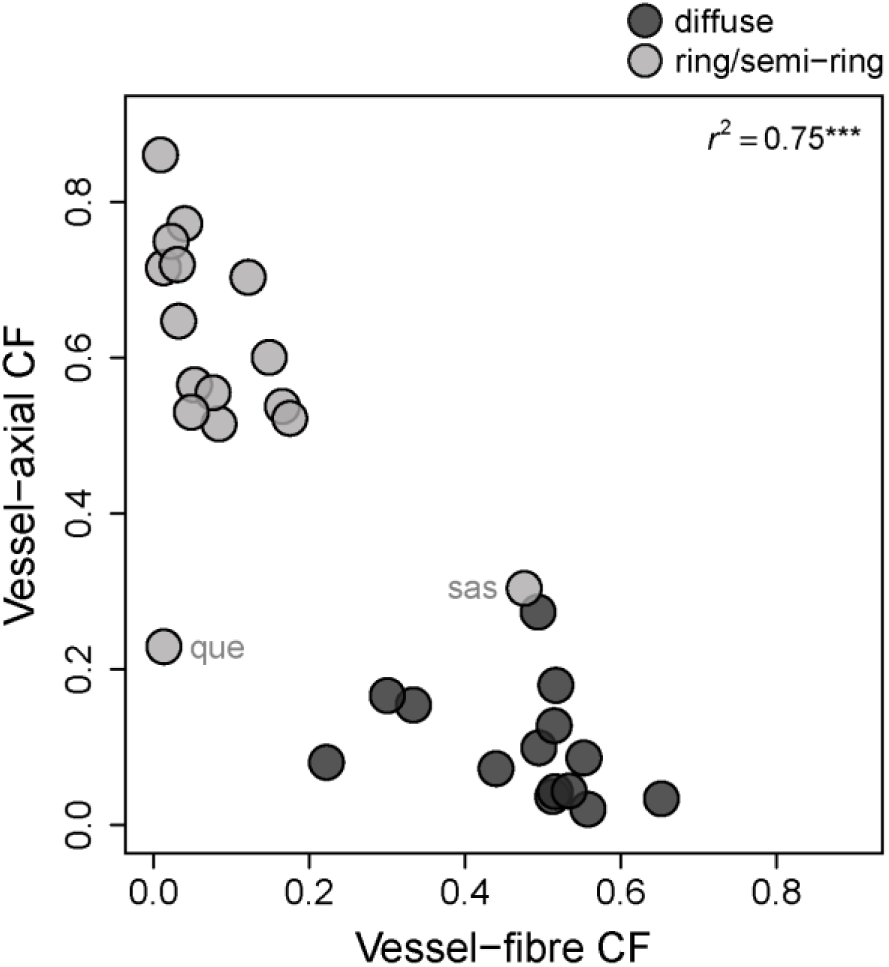
Relationship between vessel-axial parenchyma contact fraction and vessel-fibre contact fraction. CF: contact fraction. Legend displays porosity type. *R*^2^ is across all species. Outliers are *Quercus muehlenbergii* (que) and *Sassafras albidum* (sas). Legend illustrates porosity type.

Vessel diameter was 33 ± 7 μm (mean vessel area: 1077 ± 531 μm^2^) and D_H_ was 61 ± 25 μm averaged across all species. Vessel properties differed markedly between diffuse-porous and ring/semi-ring-porous species (Fig. 3). Mean vessel area was negatively correlated with vessel number per mm^2^ (i.e., species with small vessels had many of them, log10 *r*^2^=0.84, *P*<0.001, Fig. S2). Species with smaller vessels, measured as mean vessel area or D_H_, also tended to have higher total vessel lumen fraction (log10 mean vessel area: *r*^2^=0.49, *P*<0.001, log10 D_H_: *r*^2^=0.51, *P*<0.001, Fig. S2). Vessel size was positively correlated with axial parenchyma fraction (log10 mean vessel area: *r*^2^=0.33, *P*<0.001, log10 D_H_: *r*^2^=0.56, *P*<0.001), and negatively (albeit weakly) with ray fraction (mean vessel area: *r*^2^=0.25, *P*<0.01, D_H_: *r*^2^=0.10, *P*<0.10). All of these relationships were strongly affected by porosity, with diffuse-porous species at the small vessel area end of spectrum and ring/semi-ring-porous species at the large vessel area end.

### Anatomy and wood density

Total lumen and total wall fraction were the strongest drivers of WD (negative with lumen: *r*^2^=0.66, P<0.001, positive with wall: *r*^2^=0.60, *P*<0.001, Fig. S2). Among the specific tissues, fibre lumen and fibre wall fractions were strongest drivers of WD (negative with fibre lumen: *r*^2^=0.48, *P*<0.001, positive with fibre wall: *r*^2^=0.36, *P*<0.001). While axial+ray parenchyma fraction was positively correlated with WD (all species: *r*^2^=0.17, *P*<0.05, excluding *P. tomentosa*: *r*^2^= 0.44, *P*<0.001). Vessel lumen fraction was negatively correlated with density (all species: *r*^2^=0.13, *P*<0.1, excluding *P. tomentosa*: *r*^2^=0.23, *P*<0.01,). Mean vessel area and D_H_ were not related to WD across all species (mean vessel area: *r*^2^=0.003, *P*=0.77, D_H_: *r*^2^=0.06, *P*=0.19).

### Water storage

Relative water content (RWC) was 0.65 ± 0.13 at predawn and 0.59 ± 0.12 at midday. Lumen relative water content (RWC_L_) was 0.52 ± 0.17 at predawn and 0.45 ± 0.16 at midday (Fig. 6). Lumen volumetric water content (VWC_L_) was on average 0.25 ± 0.09 at predawn, 0.21 ± 0.08 at midday and 0.47 ± 0.09 at saturation (Table S2). The relationship between VWC_L-sat_ and total lumen fraction was strong, as expected (*r*^2^=0.63, *P*<0.001, Fig. S3). The slope of that relationship was not significantly different from 1 (major axis fit, *P*=0.53) and the intercept was not significantly different from 0 (*P*=0.29) suggesting that the two measurements are directly related to each other. Wall VWC (VWC_W_) averaged 0.16 ± 0.03 and was positively correlated with total wall fraction (*r*^2^=0.62, *P*<0.001).

**Figure 6.**
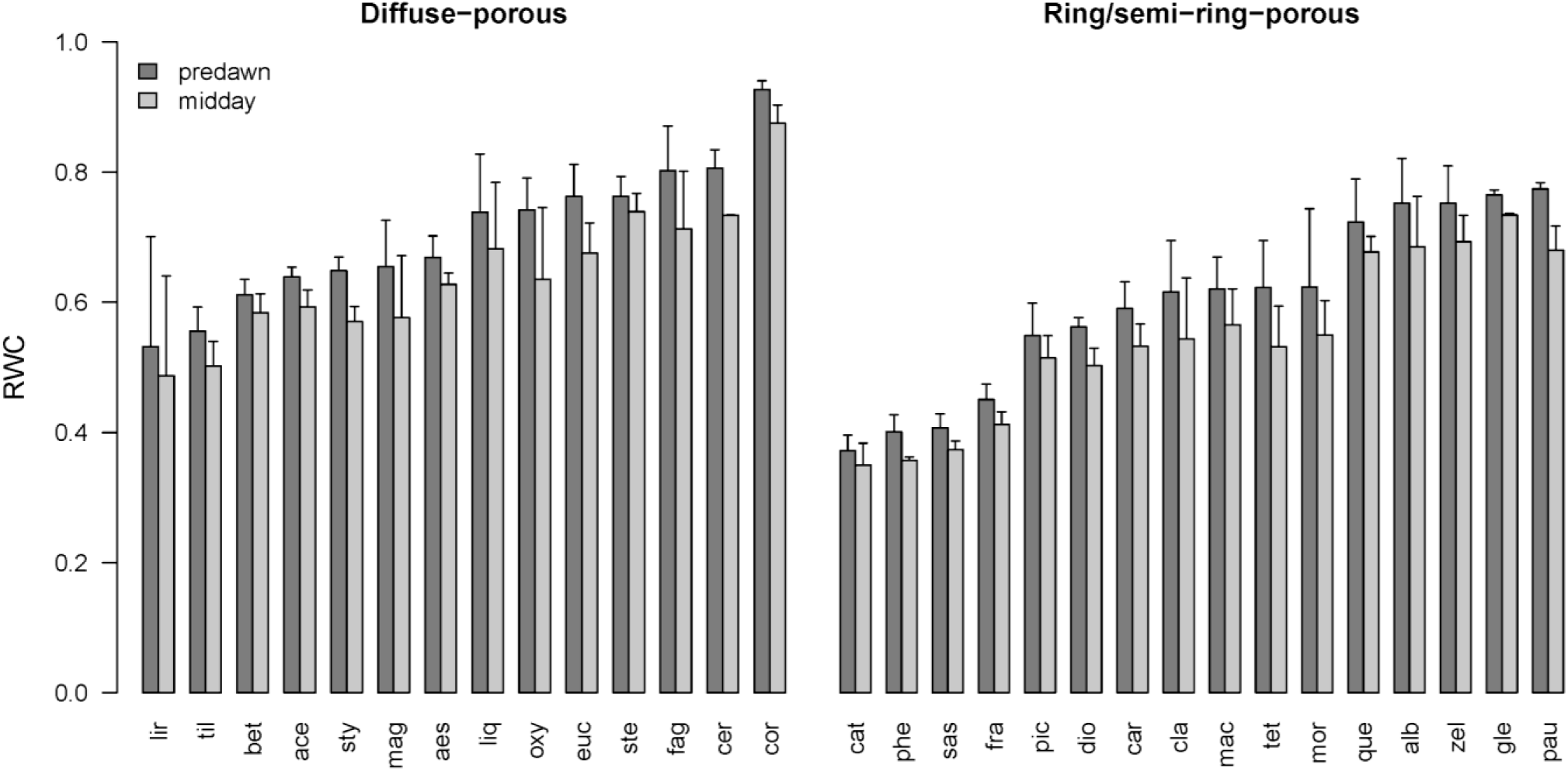
Barplot showing predawn and midday relative water content (RWC) ± 1 SD bars in diffuse-porous (a) and ring/semi-ring-porous species (b). Three-letter codes denote species (Table 1).

Predawn water content was strongly associated with midday water content for all water content indices (*r*^2^>0.97, *P*<0.001). Therefore, the following relationships are reported for predawn values only unless stated otherwise. VWC_L-pd_ was positively correlated with RWC_L-pd_ (*r*^2^=0.70, *P*<0.001) and RWC_pd_ (*r*^2^=0.49, *P*<0.001). RWC_pd_ and RWC_L_-_pd_ were strongly correlated with each other (*r*^2^=0.91, *P*<0.001), and, we use RWC_L-pd_ in subsequent analyses.

Relationships between anatomical traits and water content differed depending on water content indices and porosity type. Across all species, vessel lumen fraction was positively correlated with VWC_L_ (all species: *r*^2^=0.19, *P*<0.05, excluding *P. tomentosa*: *r*^2^=0.39, *P*<0.001, Fig. 7a). Other tissue lumen fractions were weakly or not correlated with VWC_L-pd_,(Fig. S2). VWC_L-pd_ tended to be higher in species with smaller vessels (log10 mean vessel area: all species: *r*^2^=0.20, *P*<0.05, excluding *P. tomentosa, r*^2^=0.44, *P*<0.001; log10 D_H_: all species: *r*^2^=0.32, *P*<0.01, excluding *P. tomentosa, r*^2^=0.48, *P*<0.001). These were the strongest pairwise relationships. VWC_L-pd_ was negatively correlated with WD (*r*^2^=0.19, *P*<0.05, Fig. S1) across all species. Within the two porosity groups, tissue lumen fractions or WD were not correlated with VWC_L-pd_ (Fig.S2).

**Figure 7.**
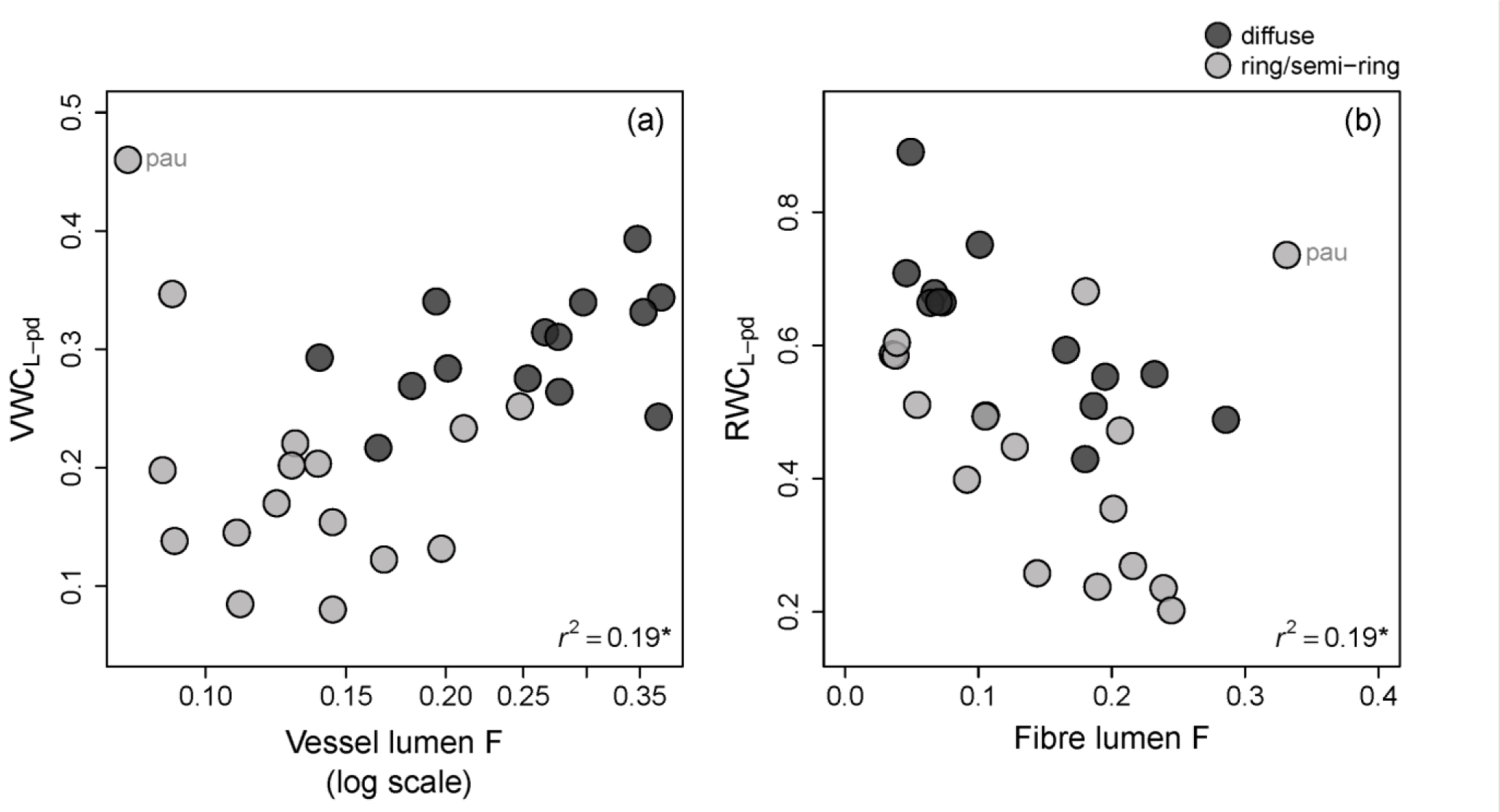
Relationships between water content and tissue lumen fractions (F): predawn lumen volumetric water content (VWC_L-pd_) vs. vessel lumen fraction (a) and predawn lumen relative water content (RWC_L-pd_) vs. fibre lumen fraction (b). Legend displays porosity type. *r*^2^ is across all species excluding *Paulownia tomentosa* (pau).

Relationships between RWC_L-pd_ and tissue fractions were stronger than for VWC_L-pd,_ and with different tissues. Across all species, the ones with higher fibre lumen fraction tended to have lower RWC_L-pd_ (all spp: *r*^2^=0.19, *P*<0.05, excluding *P. tomentosa*: *r*^2^=0.37, *P*<0.001, Fig. 7b), while other tissue lumen fractions were weakly or not related (Fig. S2). Within porosity type, fibre lumen was also the strongest, inverse correlate of RWC_L-pd_ (diffuse-porous: *r*^2^=0.65, *P*<0.001, all ring/semi-ring-porous: *r*^2^=0.07, *P*>0.1, excluding *P. tomentosa*: *r*^2^=0.58, *P*<0.01). Higher RWC_L-pd_ was associated with higher ray lumen fraction in diffuse-porous (*r*^2^=0.33, *P*<0.05) and higher axial lumen fraction in ring/semi-ring-porous (excluding *P. tomentosa*: *r*^2^=0.50, *P*<0.01). In neither of the porosity groups was vessel lumen fraction related to RWC_L-pd_.

RWC_L-pd_ was not related to WD across all species (*r*^2^=0.00, *P*=0.79), nor within ring/semi-ring-porous species (*r*^2^=0.00, *P*=0.95). But it was positively correlated within diffuse-porous species (*r*^2^=0.45, *P*=0.01).

### Day cumulative water released

The amount of water released between predawn and midday (ΔCWR_pd-md_) was on average 37 ± 14 kg per m^3^ of wood, and was positively correlated with VWC_L-pd_ (*r*^2^=0.31, *P*<0.01) and more weakly with RWC_L-pd_ (*r*^2^=0.15, *P*<0.05). ΔCWR_pd-md_ was also weakly negatively related to WD across all species (*r*^2^=15, *P*<0.05). Tissue lumen fraction or vessel-tissue contact fractions were not related to ΔCWR_pd-md_, except for a weak positive correlation with vessel lumen fraction (all species: *r*^2^=0.12, *P*<0.1, excluding *P. tomentosa*: *r*^2^=0.21, *P*<0.05) driven by porosity type.

Across diffuse-porous species, ΔCWR_pd-md_ was negatively correlated with vessel size (mean vessel area: *r*^2^=0.36, *P*<0.05, D_H_; *r*^2^=0.36, *P*<0.05) and not correlated with tissue fractions or contact fractions (on pairwise basis). Across ring/semi-ring-porous species, higher ΔCWR_pd-md_ was associated with higher VWC_L-pd_ (*r*^2=^0.55, *P*<0.01, excl. *P. tomentosa*: *r*^2^=0.39, *P*<0.05), and lower WD (*r*^2^=0.30, *P*<0.05).

### Stem water potential

Predawn stem water potential (Ψ_pd_) ranged from −0.21 to −0.93 MPa and midday stem water potential (Ψ_md_) from −0.37 to −2.1 MPa (Table S2). The largest change between predawn and midday water potential (ΔΨ_pd-md_) was 1.6 MPa and the smallest was 0.13 MPa. WD was higher in more negative Ψ_md_ (*r*^2^=0.53, *P*<0.001) and larger ΔΨ_pd-md_ (*r*^2^=0.51, *P*<0.001) species, but correlated weakly with Ψ_pd_ (*r*^2^=0.11, *P*<0.1, Fig. S2).

### Capacitance

Day capacitance was on average 53 ± 26 kg m^−3^ MPa^−1^, excluding *P. tomentosa*, and ranged from 8 to 99 kg m^−3^ MPa^−1^ (Table S2). The outlier, *P. tomentosa*, had a capacitance of 505 kg m^−3^ MPa^−1^, and was excluded from further analysis. Across all species, those with higher capacitance tended to have lower WD (*r*^2^=0.35, *P*<0.001, Fig.8a) and higher VWC_L-pd_ (*r*^2^=0.29, *P*<0.01, Fig. 8b, Table 3). Together, WD and VWC_L-pd_ explained 44% of the variation in capacitance (*r*^2^_adj_=0.44, *P*<0.001), meaning that for a given WD, species with higher VWC_L-pd_ tended to have higher capacitance and for a given VWC_L-pd,_ species with lower WD had higher capacitance. Adding tissue lumen fractions to that model did not improve its strength but adding contact fraction did (Table 3). For a given VWC_L-pd_ and WD, capacitance was higher in species with higher vessel-axial parenchyma contact fraction (*r*^2^_adj_=0.56, *P*<0.001), lower vessel-fibre contact fraction (*r*^2^_adj_=0.54, *P*<0.001) and lower vessel-ray contact fraction (*r*^2^ _adj_ =0.54, *P*<0.001, Table 3). These were the strongest models describing correlates of day capacitance across all species.

**Table 3.**
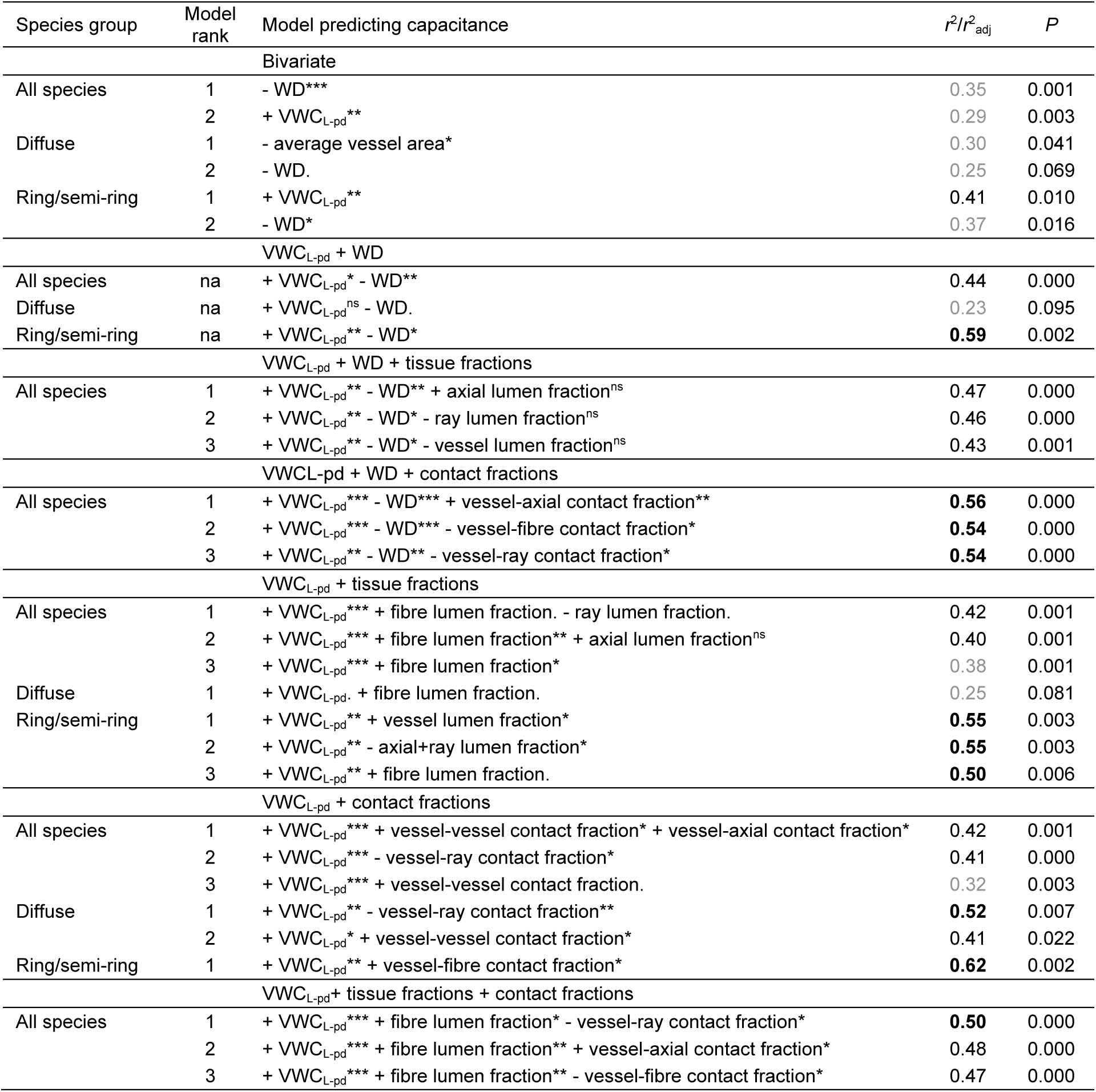
Best models predicting wood capacitance, based on *r*^2^ (for bivariate) or *r*^2^_adj_ (for multiple models). Analysed were lumen fractions of: fibre, vessel, axial parenchyma, ray parenchyma, axial+ray parenchyma, and contact fractions: vessel-fibre, vessel-vessel, vessel-axial parenchyma, vessel-ray parenchyma, vessel-axial+ray parenchyma. Three best models with *P*<0.1 are shown. For all species (n=29, excluding *Paulownia tomentosa*), we allowed maximum three explanatory variables, and for diffuse-porous (n=14) and ring/semi-ring-porous (n=15) species, we allowed maximum two explanatory variables, to avoid model overfitting. For models including vessel-fibre contact fraction among ring/semi-ring-porous species one outlier was removed (*Sassafras albidum*, see Fig. 5). VWCL-pd: lumen volumetric water content at predawn, WD: wood density, axial: axial parenchyma. Bold: *r*^2^ or *r*^2^_adj_>0.5, regular: 0.4<*r*^2^ or *r*^2^_adj_<0.5, grey: *r*^2^ or *r*^2^_adj_<0.4. Significance levels: ‘***’ *P*<0.001, ‘**’ *P*<0.01, ‘*’ *P*<0.05, ‘.’ *P*<0.1, ‘ns’ *P*>0.1.

**Figure 8.**
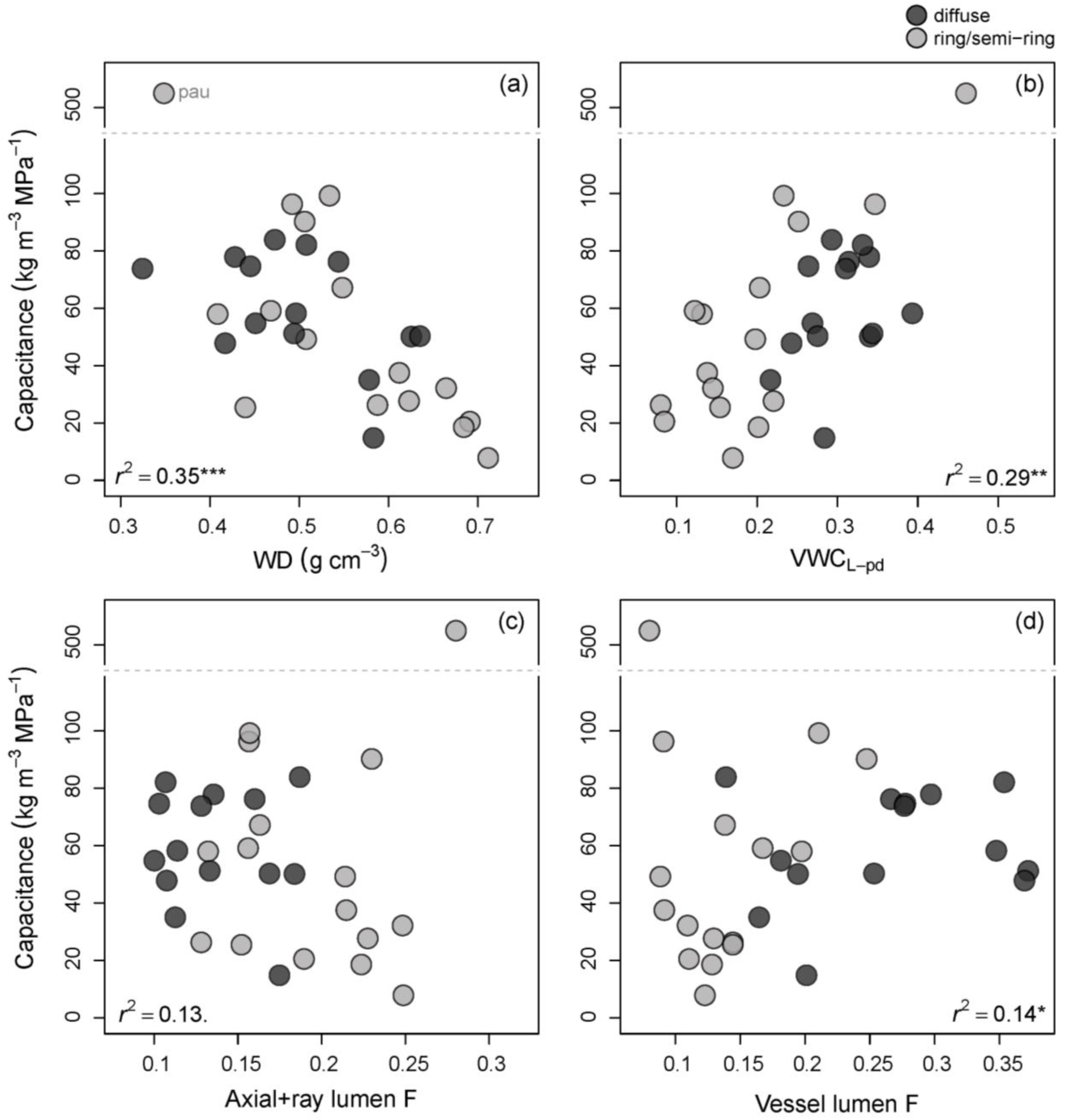
Relationships between capacitance and: wood density (WD, (a)), predawn lumen volumetric water content (VWC_L-pd_, (b)), axial+ray parenchyma lumen fraction (axial+ray lumen F, (c)), and vessel lumen fraction (vessel lumen F, (d)). Legend displays porosity type. *R*^2^ values are across all species, excluding *Paulownia tomentosa* (pau).

Relationships between capacitance and other traits differed between porosity types and across all species. Among diffuse-porous species, WD or VWC_L-pd_ were not or were only marginally correlated with capacitance (Table3). The trait combination that was most strongly related to capacitance, was VWC_L-pd_ minus vessel-ray contact fraction (*r*^2^=0.52, *P*<0.01) and plus vessel-vessel contact fraction (*r*^2^=0.41, *P*<0.05). In ring/semi-ring-porous species, VWC_L-pd_ together with WD explained the largest proportion of capacitance variation (*r*^2^_adj_=0.59, *P*<0.01, or after excluding *Sassafras albidum, r* ^2^_adj_ =0.73, *P*<0.001; *S. albidum* was the only species with the majority, if not all, fibres being gelatinous and an outlier in vessel-fibre contact fraction, Fig. 5). In addition, per given VWC_L-pd_, species with higher vessel-fibre contact fraction also had higher capacitance (*r*^2^_adj_ =0.63, *P*<0.01) but only after removing *S. albidum* (not significant, when *S. albidum* included).

When we substituted VWC_L-pd_ with RWC_pd_ in the models listed in Table 3, their strength was either comparable (‘RWC_pd_ + WD’ model across diffuse-porous species), weaker by 2-7% or not statistically significant (for other models). We also ran a set of models substituting VWC_L-pd_ with WD. Across all species, no models surpassed the ones including VWC_L-pd_ (0.28<*r*^2^_adj_ <0.37, *P*<0.01). Among diffuse-porous species, none of the models were significant (*P*>0.15), whereas across ring/semi-ring-porous species only WD was statistically significant.

Apart from the multiple regression models, species with higher capacitance tended to have less negative Ψ_pd_ (log10, *r*^2^=0.44, *P*<0.001) and narrower ΔΨ_pd-md_ (log10, *r*^2^=0.50, *P*<0.001).

## DISCUSSION

### Day capacitance and cumulative water release

Across all species, WD and VWC_L-pd_ were the strongest correlates of capacitance (Table 3, Fig. 8a, b). An inverse relationship between WD and capacitance was also found in other studies, using the bagged leaf/shoot method (Wolfe & Kursar 2015; Li *et al.* 2018), as well as in studies having used psychrometers, across a wide range of water potentials (Richards *et al.* 2014, Meinzer *et al.* 2003, 2008; Scholz *et al.* 2007; Trifilò *et al.* 2015; Jupa *et al.* 2016; Santiago *et al.* 2018; Siddiq *et al.* 2019). Taken together, these studies offer strong evidence linking capacitance to the water content and elasticity of wood tissues.

The finding that higher absolute water content (here VWC_L-pd_) conferred higher capacitance is new, yet, it makes intuitive sense, i.e., the more water there is, the more can potentially be released. Perhaps less intuitive is the result that WD was a stronger driver of capacitance than the proportions of any individual tissues. This implies that capacitance is an emergent property of the whole-wood, rather than being linked strongly to any one anatomical component. All cells in wood are tightly packed (Fig. 1), in contrast to leaves, and joined via highly lignified lamella. Therefore, any change in cell volume, resulting from water release, would require a coordinated change across neighbouring cells (Holbrook 1995). This, presumably, would be more feasible in lower WD species that have a greater whole-organ ability to shrink (Irvine & Grace 1997). This shrinkage would result from cell volume change due to water release and/or cell wall shrinkage due to adhesive forces between water and the surfaces of the water transporting conduits (Rosner, Karlsson, Konnerth & Hansmann 2009). Indeed, daily sapwood shrinkage has been observed in several angiosperm trees (Scholz *et al.* 2008; Sevanto, Hölttä & Holbrook 2011; Lintunen, Lindfors, Nikinmaa & Hölttä 2017; Hölttä *et al.* 2018), although Scholz *et al.* (2008) found no correlation between WD and diurnal sapwood shrinkage across six species. The latter study, however, encompassed a narrow WD range (∼0.42-0.62 g cm^−3^) and when lower density tissues from bark were included, strong covariation with capacitance was observed. Enlarging the number of species and broadening the wood density range in future studies would likely help to clarify this issue.

Parenchyma lumen fraction, as well as all other tissue lumen fractions, either were not correlated with capacitance or were less strongly correlated than WD. Furthermore, the tissue fractions that were significant in multiple regression models were also correlated with WD (fibre lumen and axial+ray lumen parenchyma fractions, but not vessel lumen fraction, see below). This finding is in concordance with previous studies, which also found no relationship between parenchyma fraction and capacitance in four (Jupa *et al.* 2016) and nine angiosperm species (Pratt, Jacobsen, Ewers & Davis 2007). Wood parenchyma is often assumed to be a reservoir of capacitance water (Meinzer *et al.* 2003; Steppe & Lemeur 2007; Plavcová & Jansen 2015; Vergeynst *et al.* 2015; Morris *et al.* 2016; Li *et al.* 2018; Rungwattana & Hietz 2018; Santiago *et al.* 2018) with some studies suggesting that more abundant parenchyma confers higher capacitance (Borchert & Pockman 2005; Scholz *et al.* 2011; Pratt & Jacobsen 2017; Secchi *et al.* 2017; Nardini *et al.* 2018). Here, we show that parenchyma lumen fraction does not limit capacitance (Fig. 8c). If anything, vessel lumen fraction had the strongest, albeit still weaker than WD, link to capacitance (*r*^2^=0.14, *P*<0.05, Fig. 8d) and ΔCWR_pd-md_ (*r*^2^=0.21, *P*<0.05) across all species, as well as within ring/semi-ring-porous species (multiple regression, *r*^2^_adj_=0.55, *P*<0.01, Table 3). This result is aligned with the idea that cavitating vessels might contribute to capacitance, even at xylem water potentials well above critical thresholds (Hölttä, Cochard, Nikinmaa & Mencuccini 2009; Vergeynst *et al.* 2015; Knipfer *et al.* 2019). Moreover, ΔVWC_L-pd-md_ averaged at 0.04 ± 0.02, which is small in comparison with tissue lumen fractions (Fig. 2, Table S2), suggesting that lumen fraction might not limit water release or that water could be released from multiple tissues. The interpretation of these results is further complicated by the possibility of variable water distribution across growth rings as well as within growth rings (Umebayashi *et al.* 2008, 2010). More sophisticated methods, for example, microCT or MRI (De Schepper, van Dusschoten, Copini, Jahnke & Steppe 2012; Knipfer *et al.* 2019), would be necessary to resolve these questions.

Instead of tissue lumen fractions, vessel-tissue contact fractions tended to be more strongly linked to capacitance (as independent variables in multiple regression models) across all species, as well as within porosity groups. Within ring/semi-ring-porous species, per given VWC_L-pd_, capacitance increased with higher vessel-fibre contact fraction (Table 3), suggesting that water may be released from fibres in these species (as well as vessels, see above). This is in agreement with microCT evidence of emptying fibres (and vessels) as water potential decreases (Knipfer *et al.* 2017, 2019). An interesting path forward would be to measure the size of fibre lumina and taper, because these characteristics could have an additional effect on capillary water release as capillary tension and lumen diameter are negatively correlated (Tyree & Yang 1990; Hölttä *et al.* 2009). Across diffuse-porous species, per given VWC_L-pd,_ capacitance decreased with higher vessel-ray contact fraction (Table 3). Furthermore, across all species, vessel-axial (positive), vessel-fibre (negative) and vessel-ray (negative) contact fractions were all linked to capacitance with similar strength (Table 3). Because the three contact fractions are correlated with each other (Fig. 5), it is not possible to decipher which one of these models may represent a mechanistic link. Nevertheless, the vessel-ray contact influence is consistent across all species as well as within the diffuse-porous species (Table 3). How could the inverse influence of vessel-ray contact fraction on capacitance be explained? One possibility is that, in species limited by a small volume of water released from wood (low capacitance), additional water could be supplied from bark or pith (Goldstein, Meinzer & Monasterio 1984; Cochard, Forestier & Améglio 2001; Pfautsch, Renard, Tjoelker & Salih 2015b; Pfautsch, Hölttä & Mencuccini 2015a; Mason Earles *et al.* 2016) and higher vessel-ray contact would potentially facilitate the release of this radially transported water into vessels. Partitioning capacitance between bark, wood and pith could possibly clarify the influence of contact fractions and ray parenchyma on whole-stem capacitance. We are aware of only one study (Martínez-Cabrera, Jones, Espino & Schenk 2009) that quantified contact fractions, but not in relation to any physiological trait. Our results suggest that tissue connectivity may be an important functional trait.

### Water storage

None of the studied species was fully saturated – far from it, about half of the cell lumen was empty (average RWC_L-pd_: 0.52 ± 0.17). Similar values were reported in the trunk wood of three temperate angiosperms (Longuetaud *et al.* 2016, 2017). Lack of saturation may explain why our capacitance values are lower, sometimes by an order of magnitude, than capacitance estimated as the initial slope of the water release curve (on saturated samples) in several studies of tropical (Meinzer *et al.* 2003, 2008; Carrasco *et al.* 2015; Santiago *et al.* 2018) and temperate species (Jupa *et al.* 2016), but in a similar range to Cerrado species (Scholz *et al.* 2007). Given that most of these studies measured capacitance on short xylem segments, it is possible that water may have been released from open conduits, thus resulting in an overestimation of capacitance (Tyree & Yang 1990; Jupa *et al.* 2016). Our values overlap more so with studies which used the bagged leaf method on tropical species (Zhang *et al.* 2013; Wolfe & Kursar 2015). Note that (Zhang *et al.* 2013) measured capacitance on entire stems, including bark and pith, which may explain the presence of species with high capacitance in that study. Our results also overlap with capacitance values of Australian angiosperms, estimated from excised material using psychrometers within the native operating shoot water potential range of these species (Richards *et al.* 2014), as well as with capacitance values estimated from the second, “flatter” phase of the water release curve (Jupa *et al.* 2016). Overall, these findings highlight the importance of *in natura* water content and capacitance measurements and the need for a better understanding of how these measurements compare with ones in which the water potentials were measured on small, excised samples.

WD, a direct outcome of total lumen and wall fractions, was weakly or not related to fresh water content indices, similar to other studies on three temperate angiosperms (Longuetaud *et al.* 2016, 2017) and across *ca*. 290, 180 and 100 species from Southeast Asia (Suzuki 1999; Kenzo, Tomoaki, Yuta, Joseph Jawa & Sophal 2016; Kenzo, Sano, Yoneda & Chann 2017). This lack of correlation is likely due to a considerable portion of lumen (most likely fibre lumen, Fig.7) being empty as indicated by RWC_L-pd_. These results caution against WD being taken as a direct proxy of water content in the fresh state.

VWC_L_ was more strongly correlated with capacitance than total VWC or the more commonly measured RWC, suggesting that absolute, lumen-based water indices are more relevant for capacitance than relative ones. Moreover, we estimated that the amount of wall-bound water was considerable (VWC_W-pd_ average: 0.16 ± 0.03) in comparison with lumen water content (VWC_L-pd_ average: 0.25 ± 0.09). Although, it needs to be noted, that these estimates may carry some error because of the assumption that fibre saturation point is attained at 30% moisture content in any species, which is not always the case (Kellogg & Wangaard 1969; Dlouhá *et al.* 2018). Nevertheless, these findings highlight the need to better understand the function of lumen and wall water and their explanatory power in comparison with RWC, which conflates lumen and wall water.

### Anatomical landscape

Differences between diffuse-porous and ring/semi-ring-porous species were startling not only in vessel properties, as has been well documented, but also in tissue fractions and vessel-tissue contact fractions. To our knowledge, this has not been reported before on a broad species set and/or with such detail (but see Fujiwara, Sameshima, Kuroda & Takamura 1991; Fujiwara 1992). The anatomical patterns described here agree with ones found across six angiosperm species (Jupa *et al.* 2019). Other studies also found marked differences in wood and leaf physiology between these two porosity groups (Bush *et al.* 2008; Klein 2014; von Allmen, Sperry & Bush 2015). Although this topic is beyond the scope of this study, our anatomical results raise intriguing questions regarding structure-function relationships in diffuse-porous vs. ring/semi-ring-porous species.

We found that the wall fraction per total axial or ray parenchyma was relatively high and surprisingly consistent across species (0.48 ± 0.10 in axial parenchyma and 0.36 ± 0.05 in rays, Fig.S4). Somewhat lower values for axial parenchyma were found in ten tropical species (average: 0.29, range: 0.07 to 0.53) (Ogunwusi & Ibrahim 2017). Larger values were found for rays in 50 temperate species (range 0.25-0.65; Fujiwara 1992) and similarly for ten tropical species (0.25-0.68; Ogunwusi & Ibrahim, 2017). Parenchyma walls, especially axial parenchyma, are often considered to be thin and of insignificant contribution to the overall parenchyma proportion (e.g., Martínez-Cabrera *et al.* 2009; Zheng & Martínez-Cabrera 2013; Zieminska, Butler, Gleason, Wright & Westoby 2013; Zieminska *et al.* 2015; Fortunel, Ruelle, Beauchêne, Fine & Baraloto 2014), yet the results reported here and elsewhere suggest otherwise. Moreover, parenchyma wall thickness overlaps with that of thin-wall fibres (Fujiwara *et al.* 1991; Fujiwara 1992; Jupa *et al.* 2016). As such, parenchyma walls may influence the mechanical properties of wood, across-wall water and solute transport, and result in lower than expected parenchyma lumen fraction.

### Conclusions

This work examined the anatomical correlates of wood water storage and day capacitance. Contrary to our expectations, tissue lumen fractions did not constrain capacitance. Instead, WD, VWC_L-pd_ and the connectivity between vessels and other tissues were more closely related to capacitance than were tissue lumen fractions. Our findings challenge several common assumptions: 1) that capacitance is positively related to parenchyma fraction, and 2) that wood density is negatively correlated with water content in fresh samples. Given that fresh wood was never saturated (mean RWC_pd_: 0.65 ± 0.13), we also question the functional relevance of capacitance estimated on saturated samples., at least in temperate species. Our study is limited by a lack of information on: 1) water distribution across and within growth rings, and 2) the combined and independent effects of wood, bark and pith. Addressing these issues would likely improve our understanding of capacitance and the anatomical traits that are aligned with it. Notwithstanding these limitations, our findings offer new insights into capacitance and its anatomical determinants, and suggest intriguing avenues for future research.

## ACKNOWLEDGEMENTS

KZ and ER were supported by the Arnold Arboretum of Harvard University through the Postdoctoral Putnam Fellowship and the DaRin Butz Foundation Research Internship Program, respectively. We are grateful to all the Arnold Arboretum staff, and especially: Faye Rosin (Director of Research Facilitation), Living Collections, Horticulture, and the admin staff for excellent help and support. We also would like to cordially thank the Holbrook Lab at the Department of Organismic and Evolutionary Biology of Harvard University, for discussions about capacitance and providing a pressure chamber.

## AUTHOR CONTRIBUTION

KZ designed the study and SMG and NMH contributed to concept and method development. KZ carried out fieldwork and lab measurements. KZ and ER performed anatomical analysis. KZ analysed the data and wrote first draft of the manuscript. All authors contributed to subsequent draft revisions.

## CONFLICT OF INTEREST

The authors declare no conflict of interest.

## Notes

#### Summary of Updates

Previous version had illegible forms of equations. They have now been reformatted to display correctly.

## REFERENCES

von Allmen E.I., Sperry J.S. & Bush S.E. (2015) Contrasting whole-tree water use, hydraulics, and growth in a co-dominant diffuse-porous vs. ring-porous species pair. Trees 29, 717–728.

Begg J.E. & Turner N.C. (1970) Water Potential Gradients in Field Tobacco. Plant Physiology 46, 343–346.

Blackman C.J., Pfautsch S., Choat B., Delzon S., Gleason S.M. & Duursma R.A. (2016) Toward an index of desiccation time to tree mortality under drought. Plant, Cell & Environment 39, 2342–2345.

Borchert R. & Pockman W.T. (2005) Water storage capacitance and xylem tension in isolated branches of temperate and tropical trees. Tree Physiology 25, 457–466.

Bush S.E., Pataki D.E., Hultine K.R., West A.G., Sperry J.S. & Ehleringer J.R. (2008) Wood anatomy constrains stomatal responses to atmospheric vapor pressure deficit in irrigated, urban trees. Oecologia 156, 13–20.

Carrasco L.O., Bucci S.J., Francescantonio D.D., Lezcano O.A., Campanello P.I., Scholz F.G., … Goldstein G. (2015) Water storage dynamics in the main stem of subtropical tree species differing in wood density, growth rate and life history traits. Tree Physiology 35, 354–365.

Chapotin S.M., Razanameharizaka J.H. & Holbrook N.M. (2006a) Baobab trees (Adansonia) in Madagascar use stored water to flush new leaves but not to support stomatal opening before the rainy season. New Phytologist 169, 549–559.

Chapotin S.M., Razanameharizaka J.H. & Holbrook N.M. (2006b) Water relations of baobab trees (Adansonia spp. L.) during the rainy season: does stem water buffer daily water deficits? Plant, Cell & Environment 29, 1021–1032.

Christoffersen B.O., Gloor M., Fauset S., Fyllas N.M., Galbraith D.R., Baker T.R., … Meir P. (2016) Linking hydraulic traits to tropical forest function in a size-structured and trait-driven model (TFS v.1-Hydro). Geoscientific Model Development; Katlenburg-Lindau 9, 4227–4255.

Clearwater M.J. & Meinzer F.C. (2001) Relationships between hydraulic architecture and leaf photosynthetic capacity in nitrogen-fertilized Eucalyptus grandis trees. Tree Physiology 21, 683–690.

Cochard H., Forestier S. & Améglio T. (2001) A new validation of the Scholander pressure chamber technique based on stem diameter variations. Journal of Experimental Botany 52, 1361–1365.

De Schepper V., van Dusschoten D., Copini P., Jahnke S. & Steppe K. (2012) MRI links stem water content to stem diameter variations in transpiring trees. Journal of Experimental Botany 63, 2645–2653.

Dlouhá J., Alméras T., Beauchêne J., Clair B. & Fournier M. (2018) Biophysical dependences among functional wood traits. Functional Ecology 32, 2652–2665.

Fortunel C., Ruelle J., Beauchêne J., Fine P.V.A. & Baraloto C. (2014) Wood specific gravity and anatomy of branches and roots in 113 Amazonian rainforest tree species across environmental gradients. New Phytologist 202, 79–94.

Fox J. & Weisberg S. (2011) An {R} companion to applied regression., Second. Thousand Oaks CA: Sage.

Fujiwara S. (1992) Anatomy and properties of Japanese hardwoods II. Variation of dimensions of ray cells and their relation to basic density. IAWA Bulletin n.s. 13, 397–402.

Fujiwara S., Sameshima K., Kuroda K. & Takamura N. (1991) Anatomy and properties of Japanese hardwoods. I. Variation of fibre dimensions and tissue proportions and their relation to basic density. IAWA Bulletin n.s. 12, 419–24.

Gartner B., Moore J. & Gardiner B. (2004) Gas in stems: Abundance and potential consequences for tree biomechanics. Tree physiology 24, 1239–50.

Gleason S.M., Blackman C.J., Cook A.M., Laws C.A. & Westoby M. (2014) Whole-plant capacitance, embolism resistance and slow transpiration rates all contribute to longer desiccation times in woody angiosperms from arid and wet habitats. Tree Physiology 34, 275–284.

Goldstein G., Andrade J.L., Meinzer F.C., Holbrook N.M., Cavelier J., Jackson P. & Celis A. (1998) Stem water storage and diurnal patterns of water use in tropical forest canopy trees. Plant, Cell & Environment 21, 397–406.

Goldstein G., Meinzer F. & Monasterio M. (1984) The role of capacitance in the water balance of Andean giant rosette species. Plant, Cell & Environment 7, 179–186.

Hao G.-Y., Wheeler J.K., Holbrook N.M. & Goldstein G. (2013) Investigating xylem embolism formation, refilling and water storage in tree trunks using frequency domain reflectometry. Journal of Experimental Botany.

Holbrook N.M. (1995) Stem water storage. In Plant stems: physiology and functional morphology. pp. 151–174. Academic Press.

Holbrook N.M. & Sinclair T.R. (1992) Water balance in the arborescent palm, Sabal palmetto II. Transpiration and stem water storage. Plant, Cell & Environment 15, 401–409.

Hölttä T., Carrasco M.D.R.D., Salmon Y., Aalto J., Vanhatalo A., Bäck J. & Lintunen A. (2018) Water relations in silver birch during springtime: How is sap pressurised? Plant Biology 20, 834–847.

Hölttä T., Cochard H., Nikinmaa E. & Mencuccini M. (2009) Capacitive effect of cavitation in xylem conduits: results from a dynamic model. Plant, Cell & Environment 32, 10–21.

ImageJ.

InsideWood (2004) Published on the Internet. http://insidewood.lib.ncsu.edu.

Irvine J. & Grace J. (1997) Continuous measurements of water tensions in the xylem of trees based on the elastic properties of wood. Planta 202, 455–461.

Jupa R., Doubková P. & Gloser V. (2019) Ion-mediated increases in xylem hydraulic conductivity: seasonal differences between coexisting ring- and diffuse-porous temperate tree species. Tree Physiology.

Jupa R., Plavcová L., Gloser V. & Jansen S. (2016) Linking xylem water storage with anatomical parameters in five temperate tree species. Tree Physiology 36, 756–769.

Kellogg R. & Wangaard F. (1969) Variation in the cell-wall density of wood. Wood and Fiber Science 1, 180–204.

Kenzo T., Sano M., Yoneda R. & Chann S. (2017) Comparison of Wood Density and Water Content Between Dry Evergreen and Dry Deciduous Forest Trees in Central Cambodia. Japan Agricultural Research Quarterly: JARQ 51, 363–374.

Kenzo T., Tomoaki I., Yuta I., Joseph Jawa K. & Sophal C. (2016) Wood density and water content in diverse species from lowland dipterocarp rainforest and dry dipterocarp forest. Proceedings of the symposium “Frontier in tropical forest research: progress in joint projects between the Forest Department Sarawak and the Japan Research Consortium for Tropical Forests in Sarawak,” 94–103.

Klein T. (2014) The variability of stomatal sensitivity to leaf water potential across tree species indicates a continuum between isohydric and anisohydric behaviours. Functional Ecology 28, 1313–1320.

Knipfer T., Cuneo I.F., Earles J.M., Reyes C., Brodersen C.R. & McElrone A.J. (2017) Storage Compartments for Capillary Water Rarely Refill in an Intact Woody Plant. Plant Physiology 175, 1649–1660.

Knipfer T., Reyes C., Earles J.M., Berry Z.C., Johnson D., Brodersen C.R. & McElrone A.J. (2019) Spatiotemporal coupling of vessel cavitation and discharge of stored xylem water in a tree sapling. Plant Physiology, pp.01303.2018.

Kobayashi Y. & Tanaka T. (2001) Water flow and hydraulic characteristics of Japanese red pine and oak trees. Hydrological Processes 15, 1731–1750.

Köcher P., Horna V. & Leuschner C. (2013) Stem water storage in five coexisting temperate broad-leaved tree species: significance, temporal dynamics and dependence on tree functional traits. Tree Physiology 33, 817–832.

Lachenbruch B. & McCulloh K.A. (2014) Traits, properties, and performance: how woody plants combine hydraulic and mechanical functions in a cell, tissue, or whole plant. New Phytologist 204, 747–764.

Li X., Blackman C.J., Choat B., Duursma R.A., Rymer P.D., Medlyn B.E. & Tissue D.T. (2018) Tree hydraulic traits are coordinated and strongly linked to climate-of-origin across a rainfall gradient. Plant, Cell & Environment 41, 646–660.

Lintunen A., Lindfors L., Nikinmaa E. & Hölttä T. (2017) Xylem diameter changes during osmotic stress, desiccation and freezing in Pinus sylvestris and Populus tremula. Tree Physiology 37, 491–500.

Longuetaud F., Mothe F., Fournier M., Dlouha J., Santenoise P. & Deleuze C. (2016) Within-stem maps of wood density and water content for characterization of species: a case study on three hardwood and two softwood species. Annals of Forest Science 73, 601–614.

Longuetaud F., Mothe F., Santenoise P., Diop N., Dlouha J., Fournier M. & Deleuze C. (2017) Patterns of within-stem variations in wood specific gravity and water content for five temperate tree species. Annals of Forest Science 74, 64.

Martínez-Cabrera H.I., Jones C.S., Espino S. & Schenk H.J. (2009) Wood anatomy and wood density in shrubs: responses to varying aridity along transcontinental transects. American Journal of Botany 96, 1388–1398.

Mason Earles J., Sperling O., Silva L.C.R., McElrone A.J., Brodersen C.R., North M.P. & Zwieniecki M.A. (2016) Bark water uptake promotes localized hydraulic recovery in coastal redwood crown. Plant, Cell & Environment 39, 320–328.

McCulloh K.A., Johnson D.M., Meinzer F.C., Voelker S.L., Lachenbruch B. & Domec J.-C. (2012) Hydraulic architecture of two species differing in wood density: opposing strategies in co-occurring tropical pioneer trees. Plant, Cell & Environment 35, 116–125.

Meinzer F.C., Campanello P.I., Domec J.-C., Gatti M.G., Goldstein G., Villalobos-Vega R. & Woodruff D.R. (2008) Constraints on physiological function associated with branch architecture and wood density in tropical forest trees. Tree Physiology 28, 1609–1617.

Meinzer F.C., James S.A. & Goldstein G. (2004) Dynamics of transpiration, sap flow and use of stored water in tropical forest canopy trees. Tree Physiology 24, 901–909.

Meinzer F.C., James S.A., Goldstein G. & Woodruff D. (2003) Whole-tree water transport scales with sapwood capacitance in tropical forest canopy trees. Plant, Cell & Environment 26, 1147–1155.

Meinzer F.C., Johnson D.M., Lachenbruch B., McCulloh K.A. & Woodruff D.R. (2009) Xylem hydraulic safety margins in woody plants: coordination of stomatal control of xylem tension with hydraulic capacitance. Functional Ecology 23, 922–930.

Morris H., Plavcová L., Cvecko P., Fichtler E., Gillingham M.A.F., Martínez-Cabrera H.I., … Jansen S. (2016) A global analysis of parenchyma tissue fractions in secondary xylem of seed plants. New Phytologist 209, 1553–1565.

Nardini A., Savi T., Trifilò P. & Lo Gullo M.A. (2018) Drought Stress and the Recovery from Xylem Embolism in Woody Plants. In Progress in Botany Vol. 79. Progress in Botany, (eds F.M. Cánovas, U. Lüttge & R. Matyssek), pp. 197–231. Springer International Publishing, Cham.

Ogunwusi A. a & Ibrahim H.D. (2017) Variations in Axial and Ray Parenchyma Cells in Ten Hardwood Species Growing in Nigeria. Journal of Resources Development and Management 38, 64-68–68.

Pfautsch S., Hölttä T. & Mencuccini M. (2015a) Hydraulic functioning of tree stems—fusing ray anatomy, radial transfer and capacitance. Tree Physiology 35, 706–722.

Pfautsch S., Renard J., Tjoelker M.G. & Salih A. (2015b) Phloem as Capacitor: Radial Transfer of Water into Xylem of Tree Stems Occurs via Symplastic Transport in Ray Parenchyma. Plant Physiology 167, 963–971.

Phillips N.G., Ryan M.G., Bond B.J., McDowell N.G., Hinckley T.M. & Cermák J. (2003) Reliance on stored water increases with tree size in three species in the Pacific Northwest. Tree Physiology 23, 237–245.

Plavcová L. & Jansen S. (2015) The Role of Xylem Parenchyma in the Storage and Utilization of Nonstructural Carbohydrates. In Functional and Ecological Xylem Anatomy. (ed U. Hacke), pp. 209–234. Springer International Publishing.

Pratt R.B. & Jacobsen A.L. (2017) Conflicting demands on angiosperm xylem: Tradeoffs among storage, transport and biomechanics. Plant, Cell & Environment 40, 897–913.

Pratt R.B., Jacobsen A.L., Ewers F.W. & Davis S.D. (2007) Relationships among xylem transport, biomechanics and storage in stems and roots of nine Rhamnaceae species of the California chaparral. New Phytologist 174, 787–798.

R Core Team (2018) R: A language and environment for statistical computing. R Foundation for Statistical Computing, Vienna, Austria.

Richards A.E., Wright I.J., Lenz T.I. & Zanne A.E. (2014) Sapwood capacitance is greater in evergreen sclerophyll species growing in high compared to low-rainfall environments. Functional Ecology 28, 734–744.

Rosner S., Karlsson B., Konnerth J. & Hansmann C. (2009) Shrinkage processes in standard-size Norway spruce wood specimens with different vulnerability to cavitation. Tree Physiology 29, 1419–1431.

Ross R.J. ed. (2010) Wood handbook : wood as an engineering material, Centennial. US Department of Agriculture. USDA Forest Service. Forest Products Laboratory, Madison, WI, USA.

Rungwattana K. & Hietz P. (2018) Radial variation of wood functional traits reflect size-related adaptations of tree mechanics and hydraulics. Functional Ecology 32, 260–272.

Salomón R.L., Limousin J.-M., Ourcival J.-M., Rodríguez-Calcerrada J. & Steppe K. (2017) Stem hydraulic capacitance decreases with drought stress: implications for modelling tree hydraulics in the Mediterranean oak Quercus ilex. Plant, Cell & Environment 40, 1379–1391.

Santiago L.S., De Guzman M.E., Baraloto C., Vogenberg J.E., Brodie M., Hérault B., … Bonal D. (2018) Coordination and trade-offs among hydraulic safety, efficiency and drought avoidance traits in Amazonian rainforest canopy tree species. New Phytologist, n/a-n/a.

Schneider C.A., Rasband W.S. & Eliceiri K.W. (2012) NIH Image to ImageJ: 25 years of image analysis. Nature Methods 9, 671–675.

Scholz F.C., Bucci S.J., Goldstein G., Meinzer F.C., Franco A.C. & Miralles-Wilhelm F. (2008) Temporal dynamics of stem expansion and contraction in savanna trees: withdrawal and recharge of stored water. Tree Physiology 28, 469–480.

Scholz F.G., Bucci S.J., Goldstein G., Meinzer F.C., Franco A.C. & Miralles-Wilhelm F. (2007) Biophysical properties and functional significance of stem water storage tissues in Neotropical savanna trees. Plant, Cell & Environment 30, 236–248.

Scholz F.G., Phillips N.G., Bucci S.J., Meinzer F.C. & Goldstein G. (2011) Hydraulic Capacitance: Biophysics and Functional Significance of Internal Water Sources in Relation to Tree Size. In Size- and Age-Related Changes in Tree Structure and Function. Tree Physiology, (eds F.C. Meinzer, B. Lachenbruch & T.E. Dawson), pp. 341–361. Springer Netherlands.

Secchi F., Pagliarani C. & Zwieniecki M.A. (2017) The functional role of xylem parenchyma cells and aquaporins during recovery from severe water stress. Plant, Cell & Environment 40, 858–871.

Sevanto S., Hölttä T. & Holbrook N.M. (2011) Effects of the hydraulic coupling between xylem and phloem on diurnal phloem diameter variation: Xylem-phloem hydraulic coupling and phloem diameter variation. Plant, Cell & Environment 34, 690–703.

Siddiq Z., Zhang Y.-J., Zhu S.-D. & Cao K.-F. (2019) Canopy water status and photosynthesis of tropical trees are associated with trunk sapwood hydraulic properties. Plant Physiology and Biochemistry 139, 724–730.

Sperry J.S., Nichols K.L., Sullivan J.E.M. & Eastlack S.E. (1994) Xylem embolism in ring-porous, diffuse-porous, and coniferous trees of Northern Utah and Interior Alaska. Ecology 75, 1736–1752.

Steppe K. & Lemeur R. (2007) Effects of ring-porous and diffuse-porous stem wood anatomy on the hydraulic parameters used in a water flow and storage model. Tree Physiology 27, 43–52.

Suzuki E. (1999) Diversity in specific gravity and water content of wood among Bornean tropical rainforest trees. Ecological Research 14, 211–224.

Trifilò P., Nardini A., Gullo M.A.L., Barbera P.M., Savi T. & Raimondo F. (2015) Diurnal changes in embolism rate in nine dry forest trees: relationships with species-specific xylem vulnerability, hydraulic strategy and wood traits. Tree Physiology 35, 694–705.

Tyree M.T. & Yang S. (1990) Water-storage capacity of *Thuja, Tsuga* and *Acer* stems measured by dehydration isotherms. Planta 182, 420–426.

Umebayashi T., Utsumi Y., Koga S., Inoue S., Fujikawa S., Arakawa K., … Oda K. (2008) Conducting pathways in north temperate deciduous broadleaved trees. IAWA Journal 29, 247–263.

Umebayashi T., Utsumi Y., Koga S., Inoue S., Matsumura J., Oda K., … Otsuki K. (2010) Xylem water-conducting patterns of 34 broadleaved evergreen trees in southern Japan. Trees 24, 571–583.

Vergeynst L.L., Dierick M., Bogaerts J.A.N., Cnudde V. & Steppe K. (2015) Cavitation: a blessing in disguise? New method to establish vulnerability curves and assess hydraulic capacitance of woody tissues. Tree Physiology 35, 400–409.

Warton D.I., Duursma R.A., Falster D.S. & Taskinen S. (2011) smatr 3–an R package for estimation and inference about allometric lines. Methods in Ecology and Evolution.

Wei T. & Simko V. (2017) R package “corrplot”: visualization of a corelation matrix (Version 0.84).

Wheeler E.A. (2011) InsideWood – a web resource for hardwood anatomy. IAWA Journal 32, 199–211.

Wolfe B.T. & Kursar T.A. (2015) Diverse patterns of stored water use among saplings in seasonally dry tropical forests. Oecologia 179, 925–936.

Zanne A.E., Westoby M., Falster D.S., Ackerly D.D., Loarie S.R., Arnold S.E.J. & Coomes D.A. (2010) Angiosperm wood structure: global patterns in vessel anatomy and their relation to wood density and potential conductivity. American Journal of Botany 97, 207–215.

Zhang Y.-J., Meinzer F.C., Qi J.-H., Goldstein G. & Cao K.-F. (2013) Midday stomatal conductance is more related to stem rather than leaf water status in subtropical deciduous and evergreen broadleaf trees. Plant, Cell & Environment 36, 149–158.

Zheng J. & Martínez-Cabrera H.I. (2013) Wood anatomical correlates with theoretical conductivity and wood density across China: evolutionary evidence of the functional differentiation of axial and radial parenchyma. Annals of Botany 112, 927–935.

Ziemińska K., Butler D.W., Gleason S.M., Wright I.J. & Westoby M. (2013) Fibre wall and lumen fractions drive wood density variation across 24 Australian angiosperms. AoB PLANTS 5.

Ziemińska K., Westoby M. & Wright I.J. (2015) Broad Anatomical Variation within a Narrow Wood Density Range—A Study of Twig Wood across 69 Australian Angiosperms. PLoS ONE 10, e0124892.

Zimmermann M.H. (1983) Xylem Structure and the Ascent of Sap. Springer-Verlag.

