## Supplemental Figures for "Wood capacitance is related to water content, wood density, and anatomy across 30 temperate tree species"

Figure S1. Relationship between two measures of capacitance. 'Capacitance' (presented in the Results) was estimated as the midday to predawn difference in cumulative water released per sample volume divided by the difference in stem water potential. 'Capacitance$_{F-D}$' was estimated as the predawn to midday difference in sample water content per sample volume divided by the difference in stem water potential. See Materials and Methods in the main text for the equations.

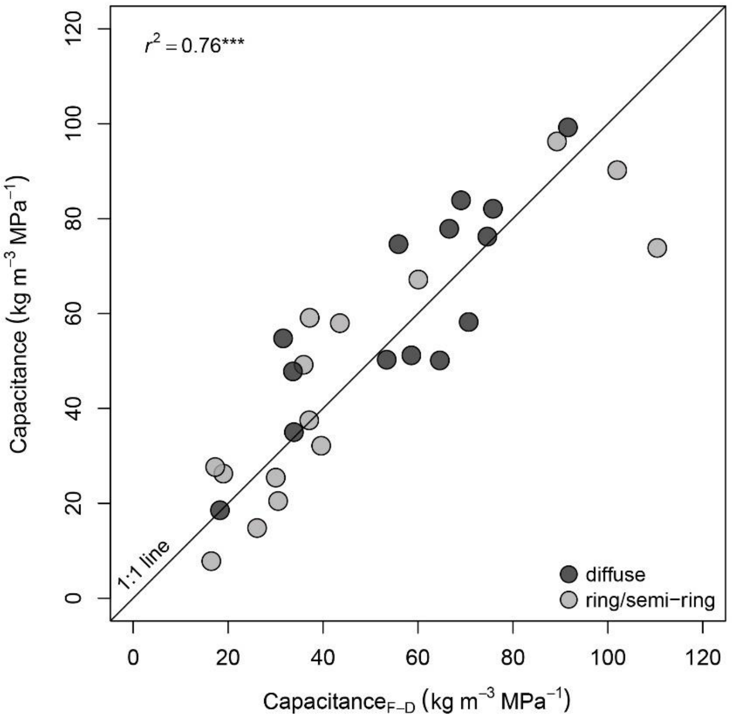

Figure S2. Scatterplot matrix of selected traits. *Paulownia tomentosa* is excluded from capacitance plots. Grey symbols: ring/semi-ring-porous species, black symbols: diffuse-porous species. Each symbol corresponds to species average (n=3).

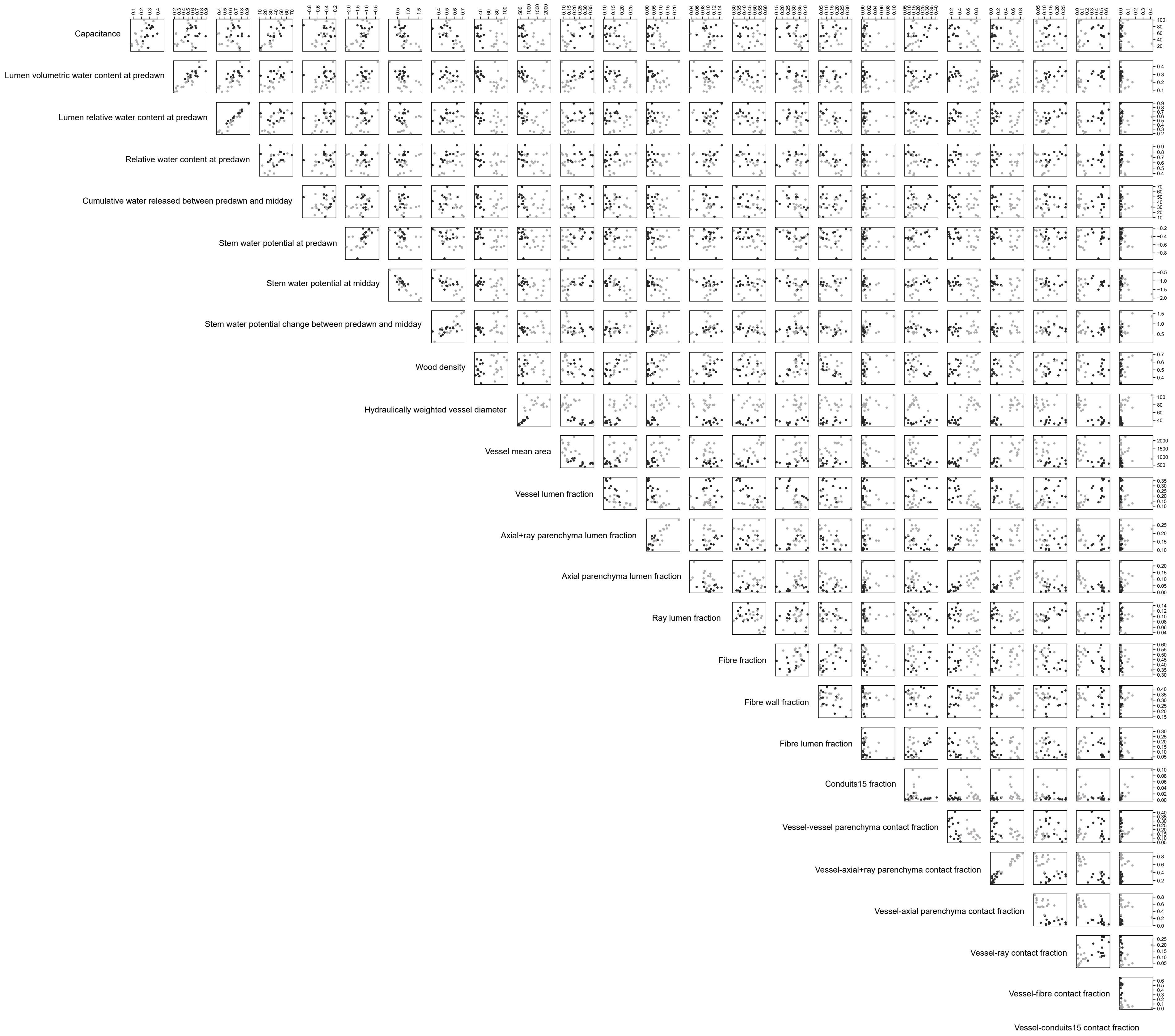

Figure S3. Relationship between total lumen fraction and volumetric water content (VWC) indices: saturated ($VWC_{sat}$), lumen saturated ($VWC_{L\text{-}sat}$) and lumen at predawn ($VWC_{L\text{-}pd}$).

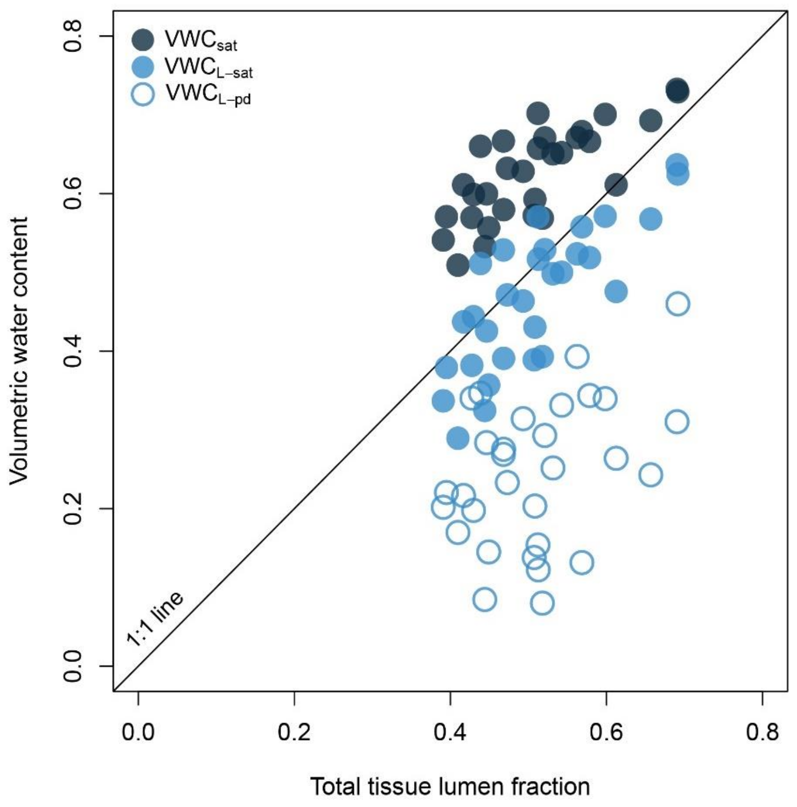

Figure S4. A schematic illustrating cross-sections of a hypothetical cell with wall area (W) to total cell area (W+L: wall+lumen) ratio equal to: 0.25, 0.50 and 0.75.

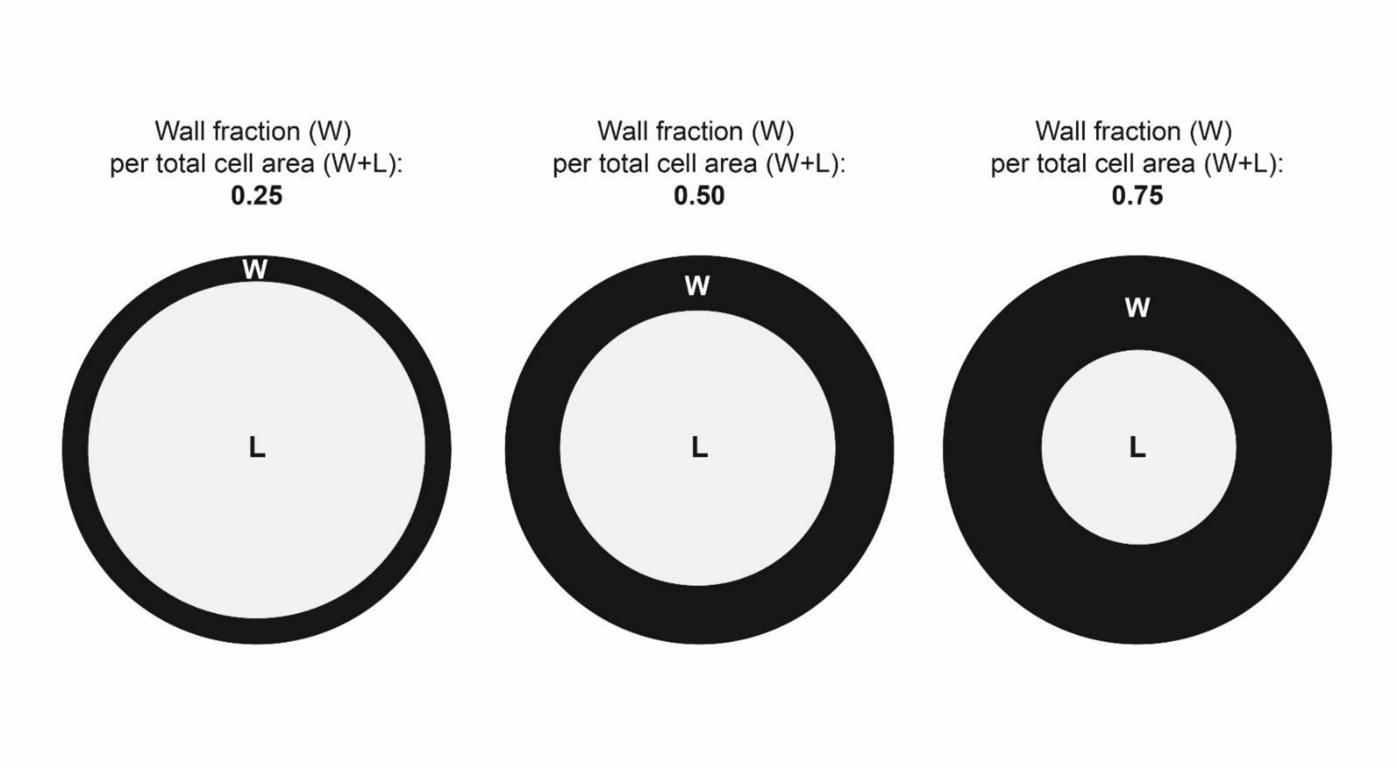
