## Supplemental Tables for "Wood capacitance is related to water content, wood density, and anatomy across 30 temperate tree species"

Table S1. List of studied species, the Arnold Arboretum accession numbers and number of individuals studied. Accession numbers which overlap between anatomical and capacitance samples are in black. Accession numbers that were sampled only for one type of measurement (anatomy or capacitance) are in grey.

| Family | Species | Porosity | Accession number - anatomy | Accession number - capacitance | N trees - anatomy | N tree - capacitance | N trees in common between the measurements |
| --- | --- | --- | --- | --- | --- | --- | --- |
| Sapindaceae | *Acer saccharum* | Diffuse | 655-93*D, 655-93*A | 655-93*D, 655-93*A, 655-93*G | 2 | 3 | 2 |
| Sapindaceae | *Aesculus turbinata* | Diffuse | 1204-77*A, 219-35*A | 1204-77*A, 219-35*A, 268-84*B | 2 | 3 | 2 |
| Betulaceae | *Betula dahurica* | Diffuse | 438-68*A, 171-79*A, 1438-77*A | 438-68*A, 171-79*A, 12793*A | 3 | 3 | 2 |
| Cercidiphyllaceae | *Cercidiphyllum japonicum* | Diffuse | 1150-67*A, 178-87*C, 13008*A | 1150-67*A, 178-87*C, 13008*A | 3 | 3 | 3 |
| Cornaceae | *Cornus kousa* | Diffuse | 524-49*C, 13123*A, 524-49*A | 524-49*C, 13123*A, 524-49*A | 3 | 3 | 3 |
| Eucommiaceae | *Eucommia ulmoides* | Diffuse | 14538*A, 112-40*A, 21931*A | 14538*A, 708-68*D, 21931*B, 112-40*B | 3 | 4 | 1 |
| Fagaceae | *Fagus grandifolia* | Diffuse | 677-2008*A, 678-2008*A, 683-2008*A | 14585*E_24, 14585*E_27, 14585*E_02 | 3 | 3 | 0 |
| Altingiaceae | *Liquidambar styraciflua* | Diffuse | 135-38*C, 1248-79*A, 135-38*B | 135-38*C, 1248-79*A, 135-38*B | 3 | 3 | 3 |
| Magnoliaceae | *Liriodendron tulipifera* | Diffuse | 143-87*B, 14992*A, 14992*D | 15330*X, 15330*AJ, 15330*AC | 3 | 3 | 0 |
| Magnoliaceae | *Magnolia cylindrica* | Diffuse | 158-83*B, 397-81*B, 158-83*C | 158-83*B, 397-81*B, 397-81*A | 3 | 3 | 2 |
| Ericaceae | *Oxydendrum arboreum* | Diffuse | *15131*C, 15131*G, 480-43*J* | *510-38*I, 20954*B, 480-43*M* | 3 | 3 | 0 |
| Theaceae | *Stewartia pseudocamellia* | Diffuse | 661-70*A, 11440*B, 405-42*A | 661-70*A, 8-43*A, 1269-83*A | 3 | 3 | 1 |
| Styracaceae | *Styrax obassia* | Diffuse | 1500-77*B, 1500-77*A, 1270-83*A | 1500-77*B, 1500-77*A, 1270-83*A | 3 | 3 | 3 |
| Tiliaceae | *Tilia japonica* | Diffuse | 147-86*B, 147-86*A, 772-53*A | 147-86*B, 147-86*A, 170-91-A | 3 | 3 | 2 |
| Fabaceae | *Albizia julibrissin* | Semi-ring | 460-67*B, 1442-77*B, 125-35*A | 460-67*B, 1442-77*B, 125-35*A | 3 | 3 | 3 |
| Bignoniaceae | *Catalpa speciosa* | Semi-ring | 927-58*B, 2776*A, 2776*B | 927-58*B, 1245-79*D, 1245-79*B | 3 | 3 | 1 |
| Fabaceae | *Cladrastis kentukea* | Semi-ring | 51-87*B, 51-87*C, 16370*A | 51-87*B, 51-87*D, 13055*B | 3 | 3 | 1 |
| Ebenaceae | *Diospyros virginiana* | Semi-ring | 801-87*A, 1392-85*B, 14513*A, 14513*B | 801-87*A, 801-87*C, 801-87*B | 4 | 3 | 1 |
| Fabaceae | *Gleditsia triacanthos* | Semi-ring | 14681*A, 13218*A, 14681-1*A | 14681*A, 1589-51*A, 141-56*A | 3 | 3 | 1 |
| Scrophulariaceae | *Paulownia tomentosa* var. 'Coreana' | Semi-ring | 1703-77*E, 730-77*D, 1703-77*B | 1703-77*E, 730-77*D | 3 | 2 | 2 |
| Juglandaceae | *Carya laciniosa* | Ring | 12898*H, 22866*A, 22866*B | 12898*H, 22866*A, 806-87*B | 3 | 3 | 2 |
| Oleaceae | *Fraxinus angustifolia* ssp. *oxycarpa* | Ring | 799-83*B, 160-90*B, 160-90*A | 799-83*B, 160-90*B, 160-90*A | 3 | 3 | 3 |
| Moraceae | *Maclura pomifera* | Ring | 471-36*B, 79-46*I, 79-46*D | 471-36*B, 79-46*I, 79-46*D | 3 | 3 | 3 |
| Moraceae | *Morus alba* | Ring | 174-41*A, 174-41*C, 174-41*B | 174-41*A, 174-41*C, 47-67*A | 3 | 3 | 2 |
| Rutaceae | *Phellodendron amurense* | Ring | 181-79*A, 590-87*B, 1769-77*A | 181-79*A, 590-87*B, 13232*B | 3 | 3 | 2 |
| Simaroubaceae | *Picrasma quassioides* | Ring | 673-70*A, 610-83*A, 610-83*B | 673-70*A, 610-83*A, 610-83*B | 3 | 3 | 3 |
| Fagaceae | *Quercus muehlenbergii* | Ring | 389-91*A, 777-90*A, 389-91*C | 389-91*A, 777-90*A, 16857*C | 3 | 3 | 2 |
| Lauraceae | *Sassafras albidum* | Ring | 22915*C, 22915*E, 22915*A | 22915*C, 22915*E, 22915*A | 3 | 3 | 3 |
| Rutaceae | *Tetradium daniellii* | Ring | 702-2008*A, 701-2008*A, 126-67*A | 702-2008*A, 701-2008*A, 126-67*A | 3 | 3 | 3 |
| Ulmaceae | *Zelkova sinica* | Ring | 288-80*A, 766-84*B | 288-80*A, 766-84*B, 766-84*E | 2 | 3 | 2 |
|  |  |  |  |  | 88 | 90 | 58 |

Table S2. Summary statistics of all traits measured: median, mean, one standard deviation (SD), minimum (Min) and maximum (Max), n-fold variation and number of species, for which given trait was observed.

| Trait | Abbr | Units | Median | Mean | SD | Min | Max | n-fold variation | Number of species |
| --- | --- | --- | --- | --- | --- | --- | --- | --- | --- |
| Day capacitance – all species | capacitance | kg m^-3^ MPa | 53 | 68 | 86 | 8 | 505 | 64.7 | 30 |
| Day capacitance – excluding P. tomentosa | capacitance | kg m^-3^ MPa | 51 | 53 | 26 | 8 | 99 | 12.7 | 29 |
| Cumulative water released – predawn | CWR_pd_ | kg m^-3^ | 224 | 223 | 91 | 42 | 426 | 10.3 | 30 |
| Cumulative water released – midday | CWR_md_ | " | 259 | 260 | 90 | 71 | 452 | 6.3 | 30 |
| Cumulative water released – difference | ΔCWR_md-pd_ | " | 36 | 37 | 14 | 11 | 69 | 6.0 | 30 |
| Stem water potantiel – predawn | Ψ_pd_ | MPa | -0.43 | -0.46 | 0.17 | -0.93 | -0.21 | 4.5 | 30 |
| Stem water potantiel – midday | Ψ_md_ | " | -1.21 | -1.22 | 0.35 | -2.00 | -0.38 | 5.3 | 30 |
| Stem water potantiel – difference | ΔΨ_pd-md_ | " | 0.74 | 0.76 | 0.28 | 0.13 | 1.59 | 12.7 | 30 |
| Volumetric water content – predawn | VWC_pd_ | Unitless | 0.40 | 0.40 | 0.08 | 0.25 | 0.56 | 2.2 | 30 |
| Volumetric water content – midday | VWC_md_ | " | 0.37 | 0.37 | 0.08 | 0.24 | 0.50 | 2.1 | 30 |
| Volumetric water content – difference | ΔVWC_pd-md_ | " | 0.03 | 0.04 | 0.02 | 0.01 | 0.08 | 9.2 | 30 |
| Volumetric water content – saturated | VWC_sat_ | " | 0.63 | 0.63 | 0.06 | 0.52 | 0.74 | 1.4 | 30 |
| Lumen volumetric water content – predawn | VWC_L-pd_ | " | 0.25 | 0.25 | 0.09 | 0.08 | 0.46 | 5.7 | 30 |
| Lumen volumetric water content – midday | VWC_L-md_ | " | 0.21 | 0.21 | 0.08 | 0.06 | 0.38 | 6.4 | 30 |
| Lumen volumetric water content – difference | ΔVWC_L-pd-md_ | " | 0.03 | 0.04 | 0.02 | 0.01 | 0.08 | 14.4 | 30 |
| Lumen volumetric water content – saturated | VWC_L-sat_ | " | 0.48 | 0.47 | 0.09 | 0.30 | 0.65 | 2.1 | 30 |
| Wall volumetric water content – predawn | VWC_W-pd_ | " | 0.16 | 0.16 | 0.03 | 0.10 | 0.22 | 2.3 | 30 |
| Wall volumetric water content – midday | VWC_W-md_ | " | 0.15 | 0.16 | 0.03 | 0.10 | 0.22 | 2.3 | 30 |
| Relative water content – predawn | RWC_pd_ | " | 0.64 | 0.65 | 0.13 | 0.37 | 0.93 | 2.5 | 30 |
| Relative water content – midday | RWC_md_ | " | 0.58 | 0.59 | 0.12 | 0.35 | 0.88 | 2.5 | 30 |
| Relative water content – difference | ΔRWC_pd-md_ | " | 0.06 | 0.06 | 0.02 | 0.02 | 0.11 | 4.8 | 30 |
| Lumen relative water content – predawn | RWC_L-pd_ | " | 0.53 | 0.52 | 0.17 | 0.20 | 0.89 | 4.4 | 30 |
| Lumen relative water content – midday | RWC_L-md_ | " | 0.46 | 0.45 | 0.16 | 0.15 | 0.81 | 5.5 | 30 |
| Lumen relative water content – difference | ΔRWC_L-pd-md_ | " | 0.08 | 0.08 | 0.03 | 0.03 | 0.14 | 5.0 | 30 |
| Wood density | WD | g cm^-3^ | 0.51 | 0.53 | 0.10 | 0.32 | 0.71 | 2.2 | 30 |
| Total wall fraction |  | Unitless fraction | 0.49 | 0.48 | 0.07 | 0.30 | 0.58 | 1.9 | 30 |
| Total lumen fraction |  | " | 0.51 | 0.50 | 0.08 | 0.39 | 0.69 | 1.8 | 30 |
| Fibre fraction – wall+lumen |  | " | 0.45 | 0.45 | 0.08 | 0.30 | 0.60 | 2.0 | 30 |
| Fibre fraction – lumen |  | " | 0.14 | 0.14 | 0.08 | 0.04 | 0.33 | 9.3 | 30 |
| Fibre fraction – wall |  | " | 0.32 | 0.31 | 0.07 | 0.15 | 0.42 | 2.8 | 30 |
| Axial+ray parenchyma fraction – wall+lumen |  | " | 0.28 | 0.28 | 0.08 | 0.16 | 0.43 | 2.7 | 30 |
| Axial+ray parenchyma fraction – lumen |  | " | 0.16 | 0.17 | 0.05 | 0.10 | 0.28 | 2.8 | 30 |
| Axial+ray parenchyma fraction – wall |  | " | 0.11 | 0.11 | 0.04 | 0.05 | 0.18 | 3.9 | 30 |
| Axial parenchyma fraction – wall+lumen |  | " | 0.10 | 0.13 | 0.09 | 0.01 | 0.30 | 40.6 | 30 |
| Axial parenchyma fraction – lumen |  | " | 0.06 | 0.07 | 0.05 | 0.01 | 0.23 | 42.1 | 30 |
| Axial parenchyma fraction – wall |  | " | 0.05 | 0.06 | 0.04 | 0.00 | 0.13 | 72.1 | 30 |
| Ray fraction – wall+lumen |  | " | 0.15 | 0.15 | 0.04 | 0.06 | 0.25 | 3.9 | 30 |
| Ray fraction – lumen |  | " | 0.10 | 0.10 | 0.03 | 0.04 | 0.15 | 3.9 | 30 |
| Ray fraction – wall |  | " | 0.06 | 0.06 | 0.02 | 0.02 | 0.10 | 5.3 | 30 |
| Vessel fraction – wall+lumen |  | " | 0.22 | 0.24 | 0.11 | 0.09 | 0.45 | 4.9 | 30 |
| Vessel fraction – lumen |  | " | 0.17 | 0.20 | 0.09 | 0.08 | 0.37 | 4.7 | 30 |
| Vessel fraction – wall |  | " | 0.04 | 0.05 | 0.02 | 0.01 | 0.10 | 8.0 | 30 |
| Living fibre fraction – wall+lumen |  | " | 0.006 | 0.026 | 0.037 | 0.001 | 0.105 | 161.2 | 8 |
| Living fibre fraction – lumen |  | " | 0.006 | 0.011 | 0.013 | 0.001 | 0.034 | 48.7 | 8 |
| Living fibre fraction – wall |  | " | 0.008 | 0.020 | 0.026 | 0.001 | 0.071 | 109.0 | 8 |
| Conduits_15_ fraction |  | " | 0.008 | 0.016 | 0.023 | 0.001 | 0.099 | 100.0 | 29 |
| Vessel-fibre contact fraction |  | " | 0.20 | 0.27 | 0.22 | 0.01 | 0.65 | 71.1 | 30 |
| Vessel-axial+ray parenchyma contact fraction |  | " | 0.43 | 0.49 | 0.24 | 0.13 | 0.88 | 6.7 | 30 |
| Vessel-axial parenchyma contact fraction |  | " | 0.29 | 0.36 | 0.28 | 0.02 | 0.86 | 42.2 | 30 |
| Vessel-ray contact fraction |  | " | 0.11 | 0.13 | 0.07 | 0.02 | 0.27 | 11.9 | 30 |
| Vessel-vessel contact fraction |  | " | 0.15 | 0.18 | 0.09 | 0.06 | 0.41 | 6.8 | 30 |
| Vessel-conduits_15_ contact fraction |  | " | 0.01 | 0.04 | 0.08 | 0.00 | 0.42 | NA | 29 |
| Vessel mean area |  | µm^2^ | 894 | 1077 | 531 | 409 | 2230 | 5.5 | 30 |
| Vessel mean diameter |  | µm | 31 | 33 | 7 | 24 | 48 | 2.1 | 30 |
| Hydraulically weighted vessel diameter | D_H_ | " | 66 | 61 | 25 | 28 | 105 | 3.8 | 30 |
| Vessel number per area |  | mm^-2^ | 178 | 247 | 198 | 37 | 625 | 16.9 | 30 |
| Vessel size to number ratio |  | mm^2^ mm^-2^ | 0.00 | 0.00 | 0.00 | 0.00 | 0.00 | 89.9 | 30 |
| Lumen fraction in fibres | lumen fraction_fibre_ | " | 0.28 | 0.30 | 0.15 | 0.08 | 0.65 | 8.0 | 30 |
| Lumen fraction in axial+ray parenchyma | lumen fraction_axial+ray_ | " | 0.58 | 0.60 | 0.05 | 0.53 | 0.77 | 1.4 | 30 |
| Lumen fraction in axial parenchyma | lumen fraction_axial_ | " | 0.52 | 0.52 | 0.10 | 0.27 | 0.78 | 2.9 | 30 |
| Lumen fraction in rays | lumen fraction_ray_ | " | 0.62 | 0.64 | 0.05 | 0.56 | 0.74 | 1.3 | 30 |
| Lumen fraction in vessels | lumen fraction_vessel_ | " | 0.81 | 0.82 | 0.04 | 0.71 | 0.88 | 1.2 | 30 |
| Lumen fraction in living fibres | lumen fraction_living fibre_ | " | 0.52 | 0.53 | 0.16 | 0.33 | 0.74 | 2.3 | 8 |
